# Comparative genomics support reduced-genome *Paraburkholderia* symbionts of *Dictyostelium discoideum* amoebas are ancestrally adapted professional symbionts

**DOI:** 10.1101/2022.06.13.495842

**Authors:** Suegene Noh, Benjamin J. Capodanno, Songtao Xu, Marisa C. Hamilton, Joan E. Strassmann, David C. Queller

## Abstract

The social amoeba *Dictyostelium discoideum* is a predatory soil protist frequently used for studying host-pathogen interactions. A subset of *D. discoideum* strains isolated from soil persistently carry symbiotic *Paraburkholderia*, recently formally described as *P. agricolaris, P. bonniea*, and *P. hayleyella*. The three facultative symbiont species of *D. discoideum* present a unique opportunity to study a naturally occurring symbiosis in a laboratory model protist. In addition, there is a large difference in genome size between *P. agricolaris* (8.7 million base pairs) vs. *P. hayleyella* and *P. bonniea* (4.1 Mbp) and in GC content (62% vs. 59%). We took a comparative genomics approach and compared the three genomes of *D. discoideum*-symbionts to 12 additional *Paraburkholderia* genomes to test for genome evolution patterns that frequently accompany host adaptation. Overall, *P. agricolaris* is difficult to distinguish from other *Paraburkholderia* based on its genome size and content, but the two reduced genomes of *P. bonniea* and *P. hayleyella* display characteristics that support evolution in a host environment. In addition, all three *D. discoideum*-symbiont genomes have increased secretion system and motility genes that may mediate interactions with their host. Specifically, adjacent BurBor-like type 3 and T6SS-5-like type 6 secretion system operons shared among all three *D. discoideum*-symbiont genomes may be important for host interaction. Ultimately, our combined evidence supports that the reduced-genome *D. discoideum*-symbionts have evolved to be professional symbionts ancestrally adapted to their protist hosts.

## Introduction

The social amoeba *Dictyostelium discoideum* (Eumycetozoa; Dictyosteliales) is a predatory soil protist frequently used for studying host-pathogen interactions (Cosson & Soldati 2008; Bozzaro & Eichinger 2011; Dunn et al. 2017). It is also an emerging model for host-microbe symbiosis in the broad sense, which we define here as an intimate association between a eukaryotic host and a prokaryote symbiont that can result in positive, neutral, or negative fitness consequences in both parties involved (Tipton et al. 2019; Hentschel 2021; Drew et al. 2021). A subset of *D. discoideum* strains isolated from soil persistently carry intracellular gram-negative *Paraburkholderia* (Betaproteobacteria; Burkholderiales) (Brock et al. 2011; DiSalvo et al. 2015; Haselkorn et al. 2019). Multi-locus sequence typing analyses and whole genome phylogenies showed that these symbionts comprise 2 independent clades (Haselkorn et al. 2019; Brock et al. 2020). Subsequently, *P. agricolaris*, and the two sister species *P. bonniea*, and *P. hayleyella* were formally described as new species sufficiently different from any other previously described *Paraburkholderia* using genetic and phenotypic evidence (Brock et al. 2020).

The three *Paraburkholderia* symbionts of *D. discoideum* present a unique opportunity to study a naturally occurring symbiosis in a laboratory model protist. They additionally present opportunities for insight into the diversity of protist-prokaryote symbioses, which are understudied compared to the symbiotic relationships of multicellular eukaryotes and their microbial symbionts (Husnik et al. 2021). The association between *D. discoideum* and its *Paraburkholderia* symbionts appears to be facultative. *D. discoideum*-symbionts are able to simultaneously maintain a free-living and host-associated lifestyle (Haselkorn et al. 2019; DiSalvo et al. 2015). The fitness outcomes to host and symbiont appear to be context-dependent (Scott et al. in press), as with most facultative host-microbe symbioses (Drew et al. 2021). *D. discoideum* amoeba hosts generally suffer negative fitness consequences of association. When amoeba are infected with their *Paraburkholderia* symbionts in the lab, the hosts tend to eat less food bacteria during vegetative growth, migrate shorter distances as slugs during their multicellular social cycle, form shorter and smaller volume fruiting bodies, produce fewer spores, and carry other bacteria alive (secondary carriage) into their next vegetative growth cycle (Brock et al. 2011; DiSalvo et al. 2015; Shu, Brock, et al. 2018; Miller et al. 2020). However potentially important context-dependent fitness benefits to the host may be (1) increased availability of food bacteria in relatively inhospitable environments as a result of secondary carriage, and (2) improved competitive ability against other *D. discoideum* strains by potentially passing on symbiont infections or releasing *Paraburkholderia* secretions in a defensive manner (Brock et al. 2011, 2013). We know less about fitness outcomes of association for *Paraburkholderia* symbionts but *P. hayleyella* (though not *P. agricolaris*) reaches higher population densities in the presence of *D. discoideum* compared to on its own in soil medium (Garcia et al. 2019). While not a direct demonstration of any fitness benefits, *P. agricolaris* and *P. hayleyella* show positive chemotaxis toward *D. discoideum* supernatant (Shu, Zhang, et al. 2018).

We present a comparative genomics analysis of the three type strains of *Paraburkholderia* isolated from *D. discoideum* (*P. agricolaris* – BaQS159, *P. hayleyella* – BhQS11, and *P. bonniea* - BbQS859). We isolated all three strains from *D. discoideum* hosts collected at Mountain Lake Biological Station in Virginia, USA. Notably, there is a large difference in genome size between *P. agricolaris* (8.7 million base pairs) vs. *P. hayleyella* and *P. bonniea* (4.1 Mbp) and in GC content (62% vs. 59%) (Brock et al. 2020; Haselkorn et al. 2019). Genome reduction is a pattern associated with long-term host association in many symbiotic bacteria (Moran & Plague 2004; Moran 2002; Maurelli 2007; Bliven & Maurelli 2012; Merhej et al. 2013, 2009; Toft & Andersson 2010) including pathogenic *Burkholderia* (Nierman et al. 2004). Therefore, we investigated any significant differences in genome characteristics in the genomes of *D. discoideum* symbionts, and particularly in the two reduced genomes of *P. hayleyella* and *P. bonniea*. Based on what we know from the best studied endosymbiont genomes of multicellular animals we look for patterns that frequently accompany host adaptation, such as fewer genes in functional categories related to metabolism, DNA repair, and gene regulation (McCutcheon & Moran 2012; Andersson & Kurland 1998).

Because the ability to infect *D. discoideum* appears to be a shared derived trait among *Paraburkholderia* symbionts of *D. discoideum*, we focus several analyses on shared orthologous genes across the three genomes in comparison with other *Paraburkholderia*. Given the estimated large phylogenetic distance between the two *D. discoideum*-symbiont clades (Brock et al. 2020; Haselkorn et al. 2019), we pay particular attention to shared horizontally transferred genetic elements. Horizontal gene transfer generally contributes to an increase in prokaryote genomic repertoires but is subject to evolutionary processes including selection and drift as with the rest of the genome (Abby & Daubin 2007; Arnold et al. 2022; Brockhurst et al. 2019; Liu et al. 2004). In the context of symbiosis, key horizontally transferred genes can enable new symbiotic relationships (e.g. symbiosis islands) (Hacker & Carniel 2001). If host adaptation-induced genome reduction is ongoing, we expect symbiont genomes to show signs of instability in the form of excess nonfunctional horizontally transferred genetic elements (e.g. insertion sequence (IS) elements or pseudogenes) (Ochman & Davalos 2006). IS elements in particular connect the themes of genome reduction and horizontally transferred genetic elements. They often proliferate during earlier stages of host adaptation and enable genome rearrangements and deterioration (Losada et al. 2010; Manzano-Marín & Latorre 2016), eventually leading to the highly reduced genomes seen in obligate symbionts.

## Methods

### Paraburkholderia genome selection and gene prediction

Genome sequencing methods were described previously (Brock et al. 2020). Briefly, we prepared high-quality DNA from individual strains grown on SM/5 agar media using Qiagen Genomic tips (20/G). Two genomes (*P. agricolaris* and *P. bonniea*) were sequenced by the University of Washington PacBio Sequencing Services and *P. hayleyella* was sequenced by the Duke University Center for Genomic and Computational Biology, all on the PacBio SMRT II platform. Reads were assembled via HGAP versions 1.87 and 1.85 (Chin et al. 2013). After an initial round of annotation, we identified the chromosomal replication initiator *dnaA* sequence and *Initiator replication protein* in each assembly’s contig and re-oriented each contig from these genes using Circlator (Hunt et al. 2015). We used SMRT analysis software Quiver to repolish each assembly (Chin et al. 2013).

We chose the following *Paraburkholderia* with finished genomes for more detailed comparison: *P. fungorum* strain ATCC BAA-463 (Coenye et al. 2001), originally isolated from the fungus *Phanerochaete chrysosporium* (Seigle-Murandi et al. 1996), *P. sprentiae* strain WSM5005 (De Meyer et al. 2013) isolated from root nodules of the domesticated legume *Lebeckia ambigua, P. terrae* strain DSM17804 (Yang et al. 2006) isolated from broad-leaved forest soil, and *P. xenovorans* strain LB400 (Goris et al. 2004) isolated from polychlorinated biphenyl-contaminated soil. We refer to these four as our representative *Paraburkholderia* genomes. For broader scale analyses of molecular evolution and comparative genomics, we used 8 additional *Paraburkholderia* genomes that span the clade that includes *P. agricolaris, P. hayleyella* and *P. bonniea*. We added 4 plant-associated species genomes (*P. megapolitana* LMG23650, *P. phenoliruptrix* BR3459a, *P. phymatum* STM815, *P. phytofirmans* PsJN) and 4 free-living species genomes (*P. caledonica* PHRS4, *P. phenazinium* LMG2247, *P. sartisoli* LMG24000, and *P. terricola* mHS1) (Vandamme et al. 2007; Coenye et al. 2004; Vandamme et al. 2002; Sessitsch et al. 2005; Viallard et al. 1998; Vanlaere et al. 2008; Goris et al. 2002). All genomes were downloaded from NCBI and considered complete (Table S1). With the exception of *P. sartisoli* and *P. phenazinium*, all selected genomes are also finished.

We re-annotated each genome with Prokka v1.14.6 (Seemann 2014) using the annotation file of *Burkholderia pseudomallei* strain K96243 (downloaded from Burkholderia Genome DB version 9.1) as a source of known proteins. Next, we found putative pseudogenes in each genome using Pseudofinder v1.0 (Syberg-Olsen et al. 2021) with DIAMOND v2.0.6.144 (Buchfink et al. 2015) BLAST against the NCBI RefSeq non-redundant protein database (downloaded August 27, 2021) in Annotate mode. Genes predicted to be pseudogenes due to truncation (less than 65% of average length of similar genes by default) or fragmentation (adjacent predicted reading frames match the same known protein) were removed from further analysis.

### Whole genome alignment

The genome aligner progressiveMauve (Darling et al. 2010) identifies locally collinear blocks (LCBs), local alignments that occur in the same sequence order and orientation across multiple genomes. We used all 13 finished *Paraburkholderia* genomes (all genomes noted above except *P. sartisoli* and *P. phenazinium*) for the initial whole genome progressiveMauve alignment in Mauve v2015-02-25. Next we compared the positions and orientations of locally colinear blocks across our three *D. discoideum*-symbiont genomes and each of these against the four representative *Paraburkholderia* genomes to identify large scale synteny using hive plots (Krzywinski et al. 2012). We used ggraph v2.0.3 and igraph v1.2.11 in R v3.6.0 (R Core Team 2019) to generate the hive plots.

### Horizontally transferred genetic element detection

We used the ISFinder (Siguier et al. 2006) webserver (accessed October 27, 2021) and its nucleotide BLAST to identify putative IS elements to test for their proliferation in each genome. We identified the best hits by comparing overlapping hits by e-value and bit score. We retained hits that were at least 70% coverage of the IS element it matched in the database.

Genomic islands are clusters of genes of horizontally transferred origin and have been found in a range of sizes from as small as 5 to as large as 500 kilobases (Dobrindt et al. 2004; Langille et al. 2010; Bertelli et al. 2019). We applied IslandPath-DIMOB and SIGI-HMM as implemented via the IslandViewer 4 webserver (Bertelli et al. 2017). IslandPath-DIMOB uses sequence composition and mobility genes, while SIGI-HMM uses codon usage bias. We then used pairwise reciprocal megablast to determine whether any of the predicted genomic islands were shared among *D. discoideum*-symbiont genomes.

Lastly, we looked for individually occurring horizontally transferred genes using DarkHorse2 v2.0_rev09. DarkHorse2 compares individual genes against the NCBI NR database and detects genes with unusual distributions of hits by calculating a lineage probability index (LPI) score (Podell & Gaasterland 2007; Delaye et al. 2020). Vertically inherited genes will have a high LPI score because most high-scoring BLASTP hits will belong to close taxonomic relatives. Horizontally transferred genes are detected because high-scoring BLASTP hits will be taxonomically distant, leading to lower LPI scores. We used DIAMOND to perform BLASTP, then following suggestions from the author excluded self and sister species hits (*P. fungorum* for *P. agricolaris, P. bonniea* and *P. hayleyella* from each other), and set the global filter threshold to 0.02 to allow candidate matches to have bit scores up to 2% different from the best non-self match.

### Gene functional annotation

We performed broad scale functional annotation with COG (Clusters of Orthologous Groups) (Galperin et al. 2015; Tatusov et al. 2000), and KO (Kyoto Encyclopedia of Genes and Genomes (KEGG) Orthology) (Kanehisa, Sato, Kawashima, et al. 2016; Kanehisa et al. 2017). We assigned COG by RPS-BLAST (Altschul et al. 1997) against COG position-specific scoring matrices downloaded from the NCBI Conserved Domain Database (version July 31, 2019). We followed JGI MGAP v4 practices and used an e-value cutoff of 0.01 and query coverage of at least 70% to be considered a valid assignment (Huntemann et al. 2015). We assigned KO using the BlastKOALA webserver (http://kegg.jp/blastkoala/; accessed July 13-28, 2020) that performed BLASTP against the KEGG GENES database at the prokaryote Genus and eukaryote Family level (Kanehisa, Sato & Morishima 2016).

We compared functional genome composition in terms of numbers of genes observed in each COG category using agglomerative clustering and non-metric multidimensional scaling (NMDS). Both were implemented in R: agglomerative clustering using cluster v2.1.2, and NMDS using vegan v2.5-7. Both analyses support that the reduced genomes of *P. bonniea* and *P. hayleyella* comprise their own cluster while all other genomes clustered together. We compared genome statistics by cluster, including genome size, GC% (proportion of GC nucleotides in the genome), proportions of intact genes vs. pseudogenes. To determine which COG categories contribute to this difference, we used a binomial exactTest (McMurdie & Holmes 2014) using edgeR v3.26.8 (Robinson et al. 2010). Because the enrichment of functional categories of genes for the comparison of *P. bonniea* and *P. hayleyella* vs. other *Paraburkholderia* genomes may be due to maintenance of necessary genes despite genome size degradation, we looked at both normalized and raw count comparisons. For each COG category that was significantly differently detected between the two clusters both in the relative (post-normalization) and absolute (raw counts) sense, we investigated which specific COGs were contributing to the difference. We used KEGG Mapper (Kanehisa & Sato 2020) and its Reconstruct Pathway tool (https://www.genome.jp/kegg/tool/map_pathway.html; accessed March 14, 2022) to corroborate differences in pathway components in genomes based on KO annotations.

### Core genome molecular evolution

To determine orthologous genes shared among all examined genomes, we performed a pan-genome analysis using Roary v3.13.0 (Page et al. 2015) with a 70% identity threshold. To test hypotheses regarding changes in lineage-specific rates of molecular evolution in *Paraburkholderia* symbionts of *D. discoideum*, we used core genes detected in the Roary pan genome analysis. We used the whole genome species tree from Brock et al (2020) and dropped any additional taxa using the drop.tip() function in phytools v1.0-1 in R. This species tree was used throughout the subsequent molecular evolution analyses using PAML v4.9d (Yang 2007). Protein sequence multi-fasta files for each core gene were aligned with MUSCLE v3.8.31 (Edgar 2004), then converted into codon alignments using PAL2NAL v14 (Suyama et al. 2006). We ran a series of codeml analyses on each codon alignment with proportional branch lengths as recommended by the PAML manual.

We applied three alternative hypotheses that test whether patterns of molecular evolution were altered by a symbiotic lifestyle (“symbiotic”), association specifically with *D. discoideum* (“dicty”), or in the reduced genomes of *P. bonniea* and *P. hayleyella* (“reduced”) (Table S1). We compared each of their AIC scores to that of the null hypothesis (H0) that there should be no significant variation in molecular evolution across the species tree. The hypothesis with the smallest AIC score with at least a 1 point difference from the null hypothesis was considered the best fit. For genes that showed patterns of molecular evolution that best fit an alternative hypothesis, we used Wilcoxon signed-rank tests in R to compare estimates of omega (dN/dS) between groups of species.

### Essential amino acid biosynthetic repertoire

We used GapMind webserver (http://papers.genomics.lbl.gov/cgi-bin/gapView.cgi; accessed January 26, 2022) to evaluate any loss of essential amino acid biosynthesis pathways in each genome. GapMind detects genes involved in biosynthesis of 17 amino acids (all standard amino acids excluding alanine, aspartate, and glutamate) and chorismate based on MetaCyc pathways using a combination of sequence similarity and protein family profiles (Price et al. 2020, 2018). It can handle fusion proteins (two enzymes fused into a single protein) and split proteins (multi-domain enzyme split into up to two proteins).

### Protein secretion system repertoire and effector prediction

We used TXSScan (Abby et al. 2016) implemented in Galaxy/ Pasteur (accessed October 2, 2020) to identify protein secretion systems in the three focal and 12 additional *Paraburkholderia* genomes. TXSScan identifies protein secretion systems (Types I-VI and IX, including Type IV and Tight adherence (Tad) pili) and flagella based on 204 experimentally studied protein profiles. It also determines whether a secretion system is complete by the presence of mandatory and forbidden component genes by sub-type, and whether it is contained within a single operon (single locus) or across a few neighboring operons (multi locus). We used the genomes of *Burkholderia mallei* ATCC 23344 and *Burkholderia pseudomallei* K96243 here to serve as ground truth because their secretion systems are well studied. A small number of T6SS and one T3SS were classified as incomplete due to misidentifying secretion system component homologs (e.g. TssC as IglB). These were manually corrected and included in the analyses. We verified these manually corrected operons against secretion system databases and Burkholderia Genome DB v9.1 (Winsor et al. 2008).

We classified all T3SS and T6SS found in our 15 genomes. For T3SS, we used the T3Enc database v1.0 (Hu et al. 2017) and downloaded three representative amino acid sequences of thirteen categories of T3SS for the conserved component genes sctJ (inner membrane ring; IPR003282), sctN (ATPase; IPR005714), and sctV (export apparatus; IPR006302). We aligned protein sequences of each component gene using MUSCLE v3.8.31 (Edgar 2004) and made gene trees using the Le and Gascuel substitution model with FastTree v2.1.10 (Price et al. 2010). We estimated a species tree from these gene trees using ASTRID v2.2.1 (Vachaspati & Warnow 2015) and ASTRAL v5.7.8 (Zhang et al. 2018). We followed the same methods for T6SS using the SecReT6 database v3.0 (Li et al. 2015) and the conserved component genes tssB (sheath; COG3516), tssC (sheath; COG3517), and tssF (baseplate; COG3519).

We used VFDB (Virulence factors of Pathogenic Bacteria; accessed January 25, 2022) (Chen et al. 2005) and downloaded protein sequences of known *Bordatella* T3 Secreted Effectors and *Burkholderia* T3 and T6 Secreted Effectors. We used DIAMOND BLASTP and these proteins as query sequences against the predicted amino acid sequences of each genome. We also used the webserver BastionHub (accessed April 20, 2021) to predict secreted effectors. BastionHub (Wang et al. 2021) combines a hidden Markov model based approach and a machine learning approach. Lastly, we used effectiveELD (Jehl et al. 2011; Eichinger et al. 2016) on the effectiveDB server (accessed March 28, 2022) to find putative secreted proteins that contain a eukaryotic-like domain. We specifically looked for proteins with domains that belong to Pfam clans for Ank (ankyrin), TPR (tetratricopeptide repeat), LRR (leucine-rich repeat), Pentapeptide, F-box, and RING (including U-box). These domains were selected based on previous reports regarding large numbers of proteins containing eukaryotic domains among amoeba symbionts (Schmitz-Esser et al. 2010; Gomez-Valero & Buchrieser 2019; Schulz et al. 2016). InterProScan (Jones et al. 2014) webserver (accessed March 29, 2022) was used for additional investigation of secreted effector candidates.

### Paraburkholderia Genome Browser

We built a web Genome Browser for each *D. discoideum*-symbiont genome for convenient browsing of all annotated genomic features mentioned above. We used JBrowse v1 (Buels et al. 2016; Skinner et al. 2009). The front-end web application was developed in Centos Steam 8 version of Linux. We used NGINX Web Server v1.14.1 and Java OpenJDK v1.8.0_322. The browser is available at https://burk.colby.edu (it is currently behind a login while we build out its firewall so please contact S. Noh for access). The GitHub repositories supporting the browser are available at https://github.com/noh-lab/burk-browser and https://github.com/noh-lab/jbrowse-executables.

### Data and Code Availability

All analyses and figures found in this manuscript can be generated and recreated using input data and code available at the GitHub repository https://github.com/noh-lab/comparative-dicty-symbionts.

## Results and Discussion

### D. discoideum-symbionts represent two distinct categories, reduced vs. non-reduced size genomes

Sequencing using PacBio technology indicated that the genomes of all three *D. discoideum-*symbiont species are each comprised of 2 chromosomes, albeit resulting in different total genome sizes. The *P. agricolaris* genome was more than twice the size of both *P. bonniea* and *P. hayleyella* (8.7 vs. 4.1 million base pairs). The overall gene content (CDS) comparison was also proportionate, with approximately 7700 genes predicted for *P. agricolaris* as opposed to approximately 3600 genes for the reduced genomes (Table 1). The genome size and gene count of *P. agricolaris* is on par with other the *Paraburkholderia* genomes we examined (Table 1).

**Table 1.** Genome statistics of *Paraburkholderia* symbionts of *D. discoideum* and other representative *Paraburkholderia* strains for comparison

|  | *P. agricolaris*<br>**BaQS159** | *P. bonniea*<br>**BbQS859** | *P. hayleyella*<br>**BhQS11** |
| --- | --- | --- | --- |
| Scaffold count | 2 | 2 | 2 |
| Genome size<br>(chr1, chr2) | 8,721,420<br>(4,816,966,<br>3,904,454) | 4,098,182<br>(3,175,376,<br>922,806) | 4,125,700<br>(3,295,139,<br>830,561) |
| GC content (%) | 61.6 | 58.7 | 59.2 |
| Genes (total) | 7,811 | 3,600 | 3,686 |
| CDS (total) | 7,721 | 3,531 | 3,610 |
| Pseudogene count | 579 | 265 | 315 |
| rRNA | 18 | 12 | 12 |
| tRNA | 71 | 56 | 63 |
| Locality (host) | *D. discoideum* in Virginia, USA | *D. discoideum* in Virginia, USA | *D. discoideum* in Virginia, USA |

|  | ***P. fungorum***<br>**ATCC BAA-463** | ***P. sprentiae***<br>**WSM5005** | ***P. terrae* DSM**<br>**17804** | ***P. xenovorans***<br>**LB400** |
| --- | --- | --- | --- | --- |
| Scaffold count | 4 | 5 | 4 | 3 |
| Genome size (total) | 9,058,983 | 7,829,542 | 10,062,489 | 9,702,951 |
| GC content | 61.8 | 63.2 | 61.9 | 62.6 |
| Genes (total) | 8,260 | 7,185 | 9,045 | 8,760 |
| CDS (total) | 8,174 | 7,087 | 8,957 | 8,675 |
| Pseudogene count | 746 | 857 | 803 | 845 |
| rRNA count | 18 | 21 | 18 | 18 |
| tRNA count | 67 | 76 | 69 | 66 |
| Locality (host) | White-rot fungus *Phanerochaete chrysosporium* in Sweden | Domesticated legume *Lebeckia ambigua* in Western Australia | Broad-leaved forest soil in South Korea | PCB-contaminated soil in USA |

\*Genes and CDS counts were predicted by Prokka and are inclusive of pseudogenes
\*Pseudogenes were predicted by Pseudofinder

Whole genome alignments of all ten finished genomes found 153 locally colinear blocks, ranging in sizes as small as 262 base pairs and as large as 208,252 base pairs in the *P. agricolaris* genome. *P. bonniea* and *P. hayleyella* share with each other considerable synteny but also possess a large inverted region relative to each other (Figure 1). Both of these reduced genomes show extensive genome rearrangement compared to the genome of *P. agricolaris*, or any of the other *Paraburkholderia* genomes (Figure 1 & S1). The genome of *P. agricolaris* shares a large degree of synteny with other *Paraburkholderia* genomes in chromosome 1 as indicated by the overall lack of gaps toward the center of each hive plot (Figure S1).

**Figure 1.**
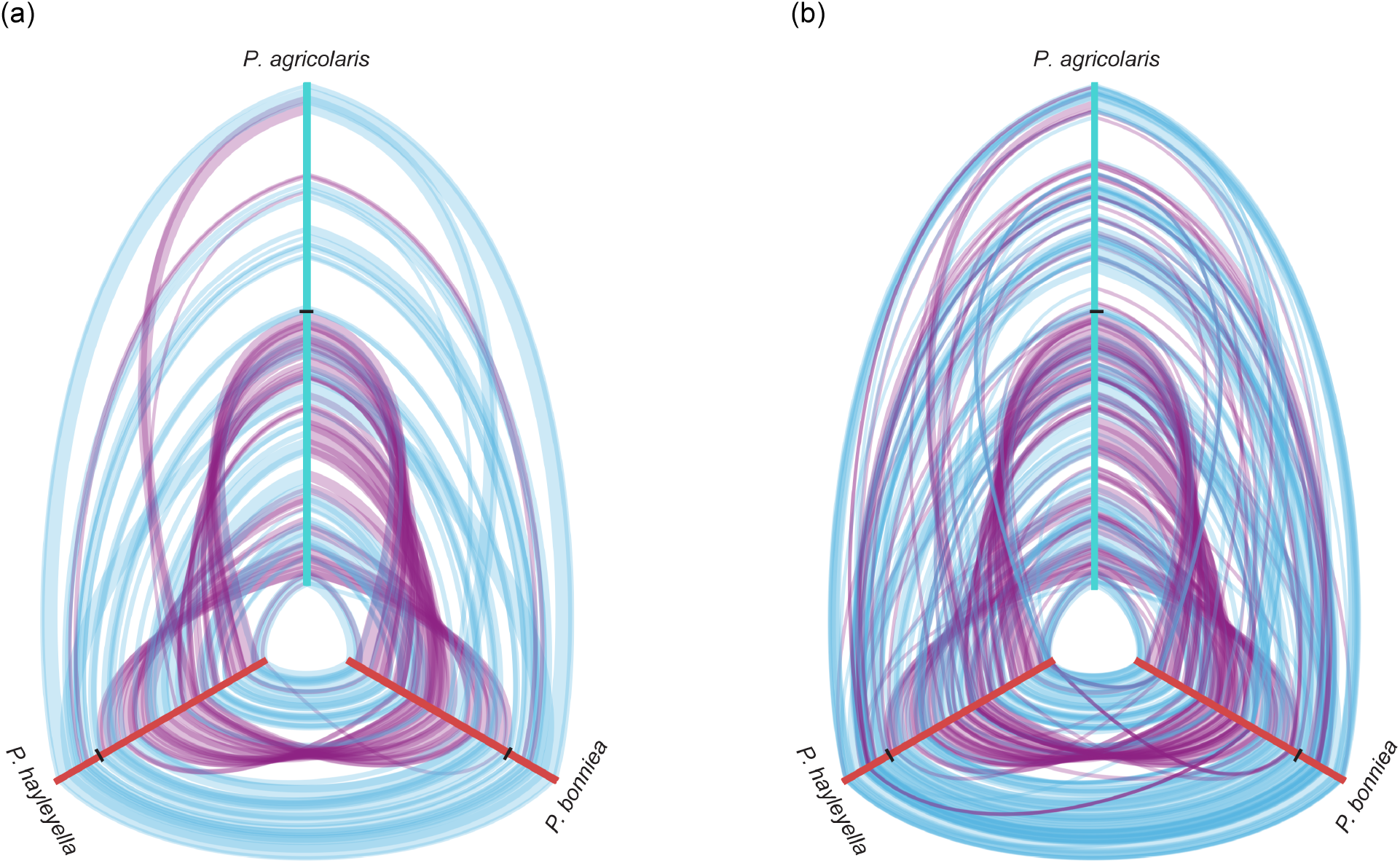
Hive plots of whole genome comparisons of *D. discoideum*-symbiont genomes. Locally colinear blocks between pairs of genomes are shown as bands that connect the axes (genomes). Only blocks above the median size are shown on the left (a) for visual clarity, while all blocks are shown on the right (b). Alignment of locally colinear blocks are distinguished between forward (blue) and reverse (purple) orientation. Axes are oriented center out, and boundaries between chromosomes are shown as ticks.

We found few IS elements in the *D. discoideum*-symbiont genomes. *P. agricolaris* has the IS elements IS1090 (6 copies) and ISBmu21 (1 copy), *P. bonniea* has ISBp1 (2 copies) and ISBuph1 (3 copies), and *P. hayleyella* has a single ISPa37 in their genomes (Table S2). Among the other *Paraburkholderia* genomes we examined, the highest number of IS elements was found in *P. xenovorans* LB400 (62 total), while others possessed intermediate numbers ranging from 5 in *P. sprentiae* to 21 in *P. fungorum*. For reference, genomes of *B. mallei* possess between 166-218 IS elements, many of which were flanking regions that were randomly lost among the examined strains in what appears to be ongoing genome reduction (Losada et al. 2010). There was also no evidence of excess pseudogenes in the reduced genomes relative to other *Paraburkholderia* genomes (Table 1). Double-strand break repair pathways (KEGG map03440) were complete in all three *D. discoideum*-symbiont genomes. As genomes with ongoing genome reduction often have numerous IS elements and pseudogenes, and incomplete double-strand break repair pathways, the combined evidence supports that all three *D. discoideum*-symbiont genomes are relatively stable, and the two reduced genomes are currently not in flux.

### The reduced D. discoideum-symbiont genomes show evidence of functional adaptation to the host environment

For each *D. discoideum*-symbiont genome, 65-68 % of genes were annotated with COG (Figure S2), and 53-61 % with KO. The agglomerative clustering and NMDS analyses of COG category representation across genomes resulted in *P. bonniea* and *P. hayleyella* clustering with each other and apart from other *Paraburkholderia* including *P. agricolaris* (Figure 2). Further investigation of specific functional differences between the two groups (reduced genomes vs. non-reduced) indicated nine COG categories that were significantly different (exactTest, FDR << 0.01). Of these, four were consistently different in both normalized and raw counts in the same direction (Figure S3). Fewer genes than expected were detected in the reduced genomes of *P. bonniea* and *P. hayleyella* for Transcription (category K), Carbohydrate transport and metabolism (G), and Inorganic ion transport and metabolism (P). More genes than expected were found in the reduced genomes for Cell motility (N).

**Figure 2.**
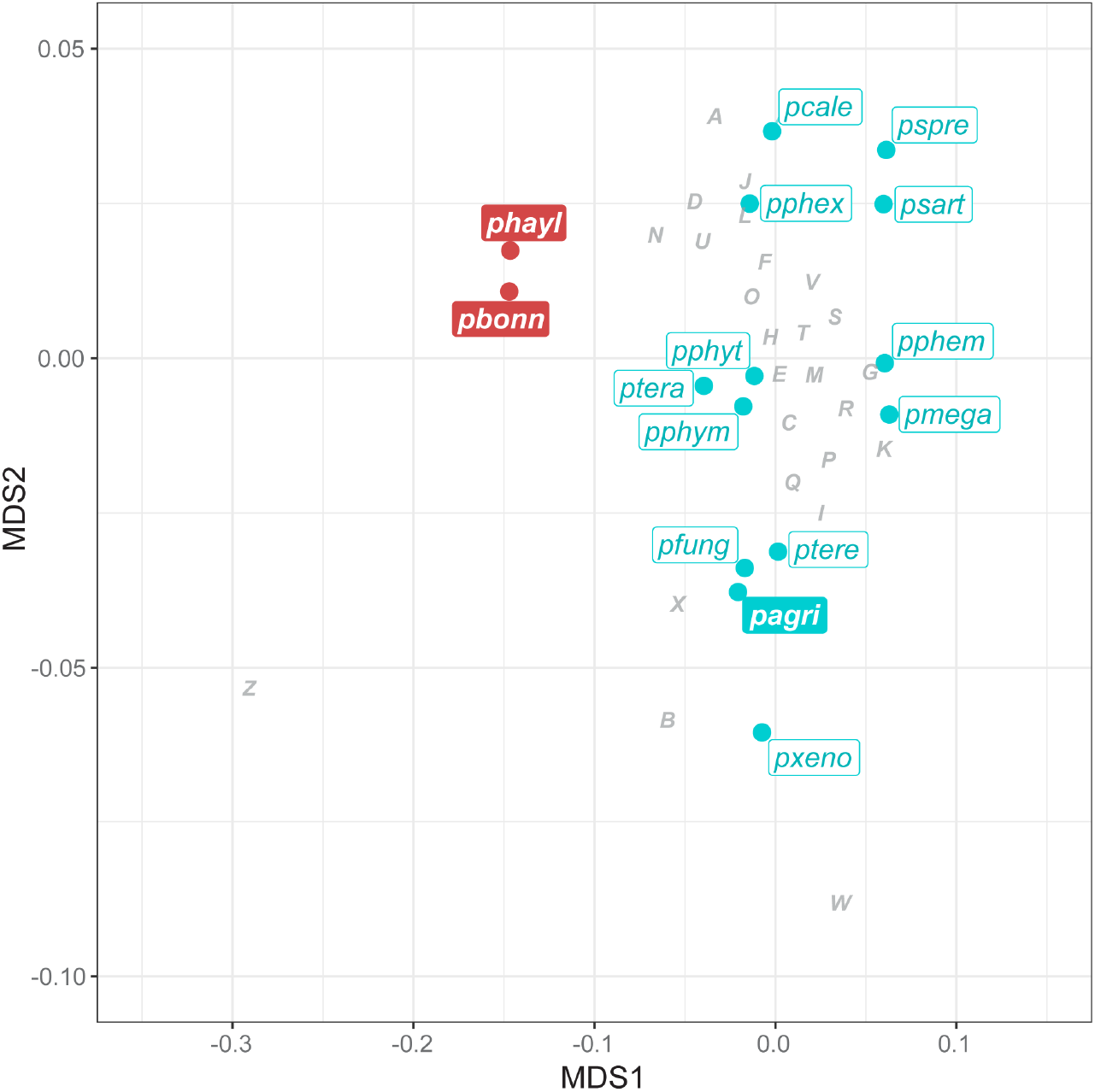
Comparison of reduced (red) and non-reduced (turquoise) genomes in terms of their functional compositions in nonmetric multidimensional space. The contributions of COG categories are projected with minor adjustments to avoid overlap with other features. (pagri = *P. agricolaris*; pbonn = *P. bonniea*; phayl = *P. hayleyella*; pcale = *P. caledonica*; pfung = *P. fungorum*, pmega = *P. megapolitana*, pphem = *P. phenazinium*; pphex = *P. phenoliruptrix*; pphym = *P. phymatum*; pphyt = *P. phytofirmans*; psart = *P. sartisoli*; pspre = *P. sprentiae*; ptera = *P. terricola*; ptere = *P. terrae*; pxeno = *P. xenovorans*) (J = Translation, ribosomal structure and biogenesis; A = RNA processing and modification; K = Transcription; L = Replication, recombination and repair; B = Chromatin structure and dynamics; D = Cell cycle control, cell division, chromosome partitioning; Y = Nuclear structure; V = Defense mechanisms; T = Signal transduction mechanisms; M = Cell wall/ membrane/ envelope biogenesis; N = Cell motility; Z = Cytoskeleton; W = Extracellular structures; U = Intracellular trafficking, secretion, and vesicular transport O = Posttranslational modification, protein turnover, chaperones; X = Mobilome: prophages, transposons; C = Energy production and conversion; G = Carbohydrate transport and metabolism; E = Amino acid transport and metabolism; F = Nucleotide transport and metabolism; H = Coenzyme transport and metabolism; I = Lipid transport and metabolism; P = Inorganic ion transport and metabolism; Q = Secondary metabolites biosynthesis, transport and catabolism; R = General function prediction only; S = Function unknown)

We looked within each COG category that was significantly different between reduced and non-reduced genomes in more detail. First, we found several flagella biosynthesis, basal body, and hook protein COGs that were more abundant in the reduced genomes than expected (Figure 3a). Flagella are often associated with bacterial virulence, not only through providing motility but also adhesion, invasion, and the secretion and regulation of virulence factors (Ottemann & Miller 1997; Duan et al. 2013). Among *Burkholderia, B. pseudomallei* flagella have been shown to be necessary for post-invasion virulence in mice (Chua et al. 2003). *B. pseudomallei* and *B. thailandensis* each have two flagellar clusters, and in *B. thailandensis* the second cryptic cluster is involved in post-invasion intracellular motility (French et al. 2011). We found a second flagellar cluster in *P. bonniea* but not in the other *D. discoideum*-symbiont genomes (see also ‘Secretion systems’ section).

**Figure 3.**
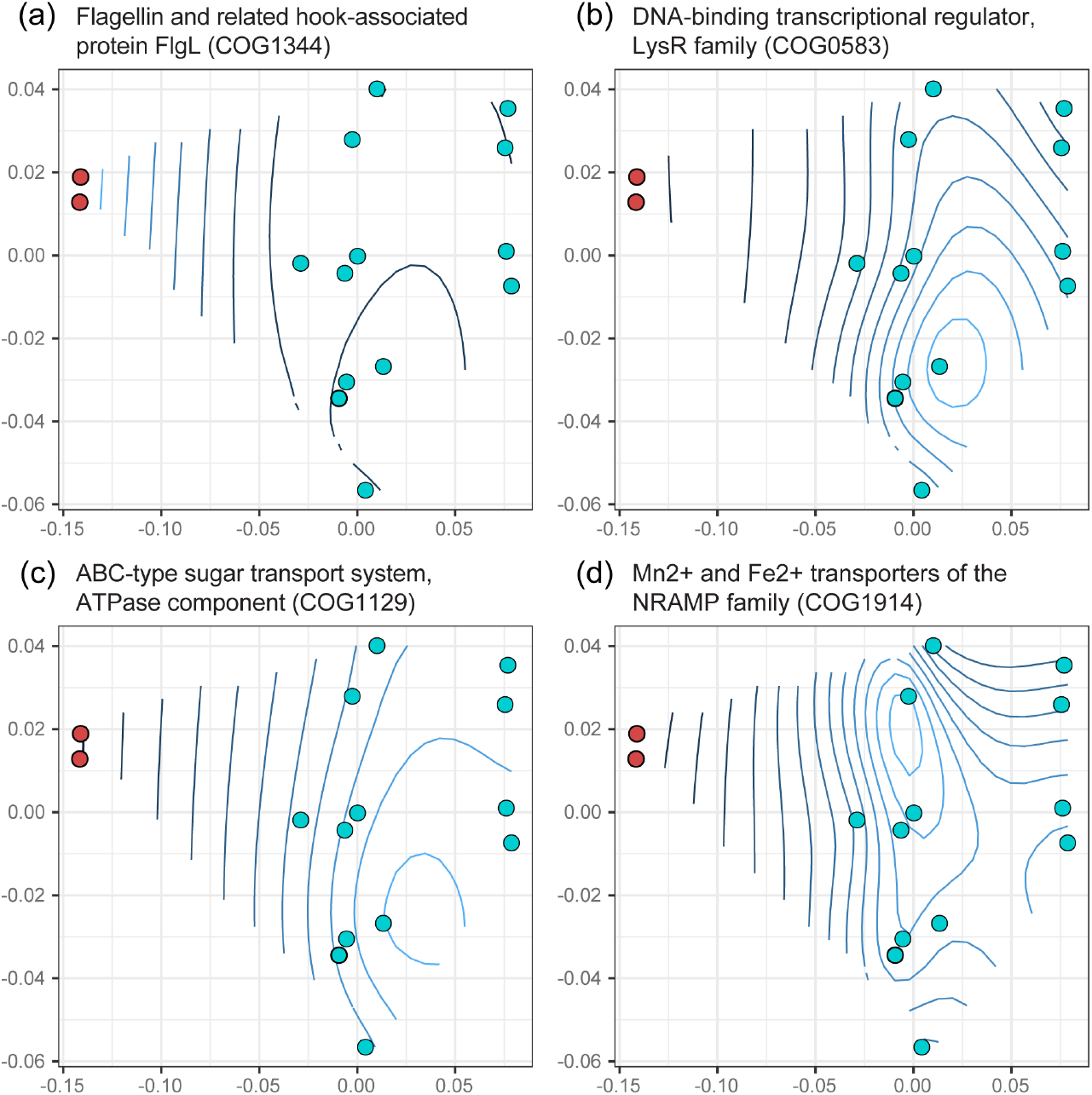
Representative individual COGs belonging to categories (a) Cell motility, (b) Transcription, (c) Carbohydrate transport and metabolism, and (d) Inorganic ion transport and metabolism (P) that were significantly overrepresented or underrepresented in the reduced genomes of *D. discoideum*-symbionts. Contours of abundances are superimposed on the nonmetric multidimensional space from Figure 2. *P. bonniea* and *P. hayleyella* are shown as red points to the left, while *P. agricolaris* is not distinguished from the other genomes in turquoise. Lighter blue contour lines indicate higher abundance compared to darker blue lines.

The other significant COG categories were less abundant in the reduced genomes than expected (Figure 3bcd). Many families of transcriptional regulator COGs were less abundant in the reduced genomes, as is often seen with reduced symbiotic bacterial genomes (Merhej et al. 2013; Wilcox et al. 2003). Similarly, several ATP binding cassette (ABC)-type sugar and metal ion transporter COGs were less abundant in the reduced genomes. ABC transporters are often reduced in number in bacteria with intracellular niches compared to extracellular or environmental ones, as intracellular environments are relatively more stable compared to extracellular environments (Garmory & Titball 2004; Harland et al. 2007).

Analysis with KEGG mapper reconstruction confirmed several missing sugar transport systems in the reduced genomes compared to *P. agricolaris*, including Sorbitol/ Mannitol, L-Arabinose, Galactofuranose, D-Xylose, Fructose, and Rhamnose. Genes encoding iron (III) transporters were also absent in the two reduced symbiont genomes compared to non-reduced *P. agricolaris*. A similar analysis of ABC transporters also revealed the presence of heme exporter proteins in both reduced genomes but not in *P. agricolaris*, and a capsular polysaccharide transport system in *P. bonniea* only. We also examined two-component system (TCS) transporters with KEGG mapper because pathogenic *Burkholderia* have multiple two-component systems related to virulence in plant and animal infection models (Schaefers 2020). Compared to *P. agricolaris*, the reduced genomes lacked genes encoding nitrate reductase proteins and chemotaxis proteins typically involved in biofilm formation through cyclic di-GMP regulation. In free-living *B. pseudomallei* these two two-component systems are linked, as the presence of nitrate has been shown to reduce intracellular cyclic di-GMP levels and inhibit biofilm formation (Mangalea et al. 2017). It appears these two-component systems and the aforementioned transporters have not been maintained under selection during host adaptation and genome reduction in *P. bonniea* and *P. hayleyella*.

### The reduced D. discoideum-symbiont genomes may experience a combination of stronger and relaxed purifying selection relative to other Paraburkholderia genomes

We identified 1673 core genes shared by the 15 *Paraburkholderia* species genomes we investigated (Figure S2). When we examined dN/dS as a signature of molecular evolution, the majority of the *Paraburkholderia* core genes showed nonsignificant variation in selection pressure across the species phylogeny. However, a large proportion of core genes (∼40 %) showed an alternative pattern of molecular evolution (Table 2). These genes show one of two patterns of molecular evolution: those that appear to experience increased selection pressure and significantly lower dN/dS once symbiotically associated with eukaryotes or specifically with *D. discoideum* (“symbiotic” and “dicty”; both Wilcoxon test P << 0.01), and those that show evidence of relaxed selection and significantly higher dN/dS in genomes of reduced size (“reduced”; Wilcoxon test P << 0.01) (Figure 4). These results indicate that the reduced genomes of *P. bonniea* and *P. hayleyella* possess a combination of genes experiencing stronger selective constraints and genes under weaker selective constraints relative to the genomes of other *Paraburkholderia*. This pattern is in contrast to genomes of obligate symbionts where the majority of genes are experiencing genetic drift and weaker selection constraints across their entire genomes (Sabater-Muñoz et al. 2017; Wernegreen 2017).

**Table 2.**
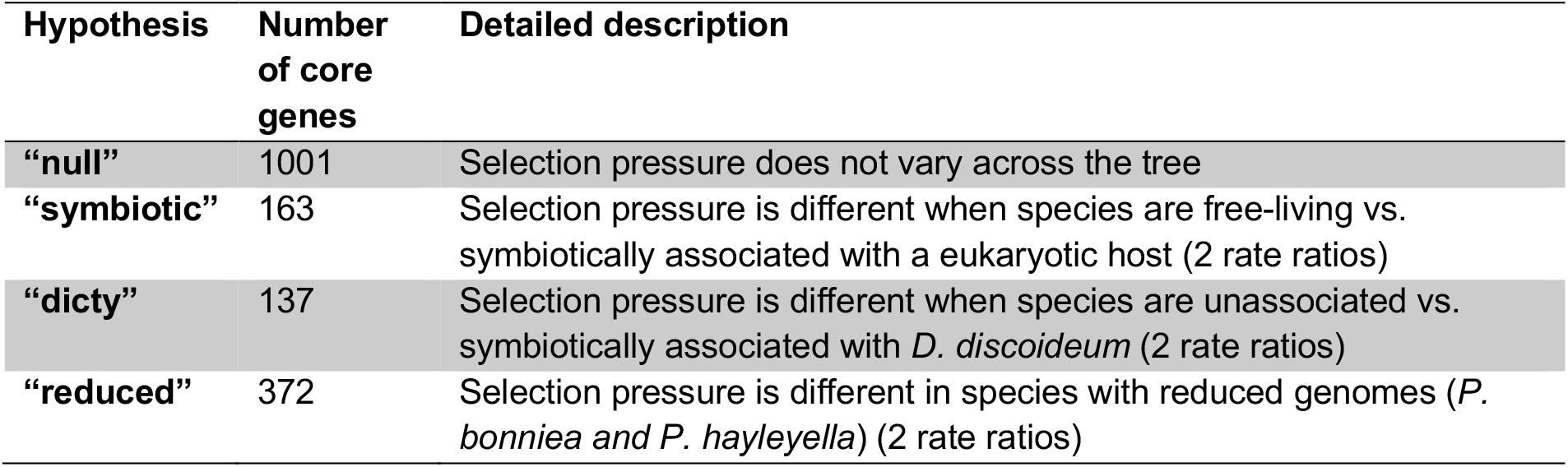
Hypotheses tested regarding molecular evolution in the 1673 core genes shared across 15 Paraburkholderia genomes.

**Figure 4.**
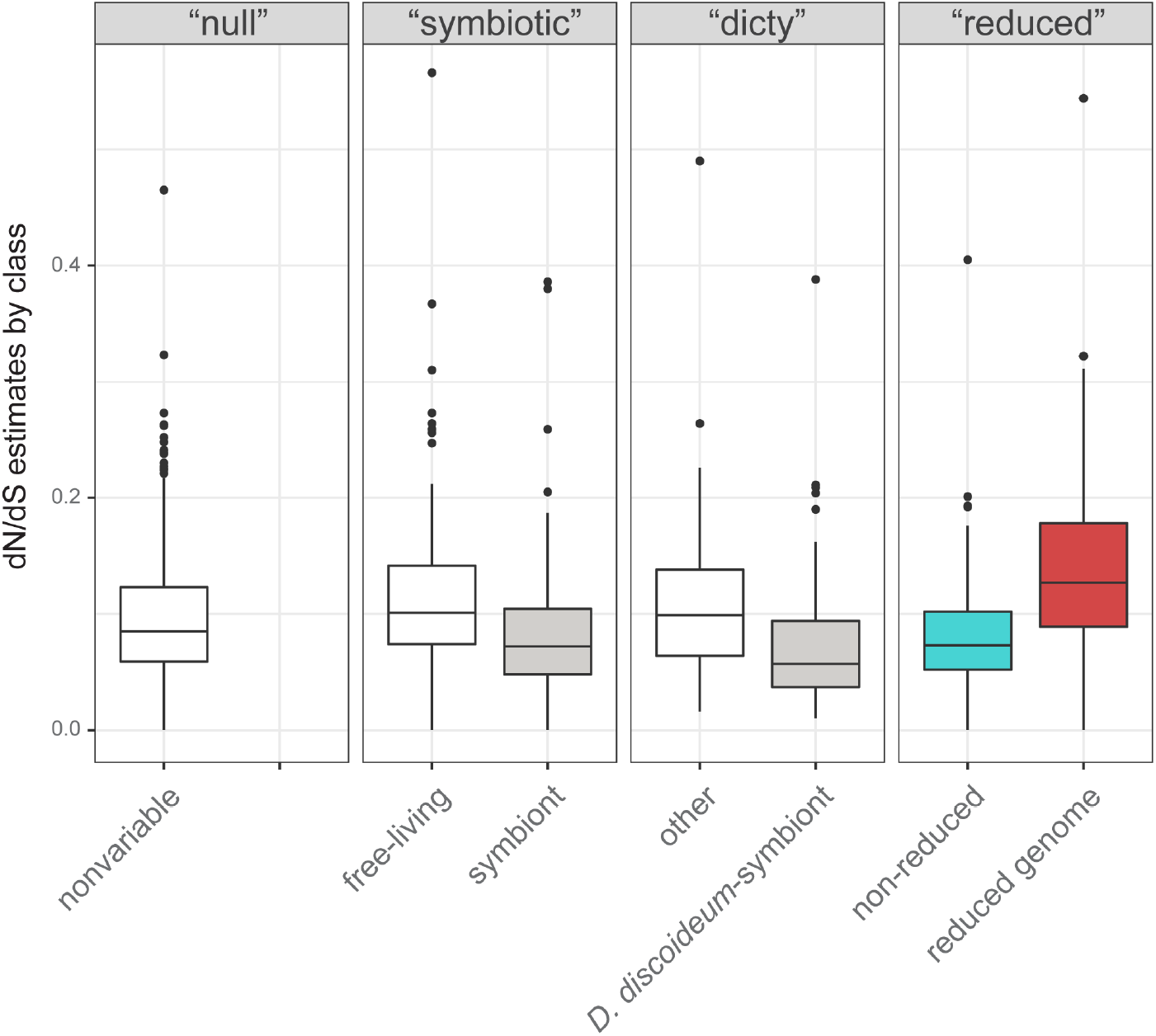
Core genes divided into the hypothesis that best predicts their patterns of molecular evolution. Core genes included genes evolving under stronger selective constraints with significantly lower dN/dS in genomes of symbionts of *D. discoideum* or other eukaryotes (“symbiotic” and “dicty”), and genes showing evidence of relaxed selective constraints with significantly higher dN/dS in the reduced genomes of *P. bonniea* and *P. hayleyella* (“reduced”).

The three *D. discoideum*-symbiont genomes shared 1977 genes total, including the 1673 core genes (Figure S2). Of the 1977 *D. discoideum*-symbiont-shared genes (inclusive of core genes), 120 were not orthologous to genes found in any of the other *Paraburkholderia* genomes we compared. These genes included type 3 and type 6 secretion system component genes (see ‘Secretion systems’ section), *bhuRSTUV* genes, and helix-turn-helix motif-containing GntR and LysR transcriptional regulators. *Bordatella* heme utilization (*bhu*) genes are virulence factors in mammalian and avian host infection (Murphy et al. 2002; Vanderpool & Armstrong 2001), and transcriptional regulators with helix-turn-helix motifs have been frequently associated with virulence in pathogens (Finlay & Falkow 1997).

### The relationship between D. discoideum and its symbionts is unlikely to be based on amino acid exchange

The gradual loss of essential amino acid biosynthetic ability is a feature of genome reduction in many microbial symbionts that have nutrient exchange relationships with their hosts (Moran et al. 2008; Lo et al. 2016; McCutcheon et al. 2019). However nutrient-dependent relationships are less likely in protist-prokaryote symbioses because protist host diets tend to be much more diverse compared to multicellular eukaryotes (Husnik et al. 2021). Accordingly, the three *D. discoideum*-symbiont species are predicted to synthesize all essential amino acids (Table S3), albeit with some variation in degrees of confidence. High confidence candidates were identified for each of the steps of amino acid biosynthesis in *P. agricolaris* but some pathways included medium confidence steps in the other two species with reduced genomes. In *P. bonniea*, the L-arginine biosynthesis pathway contained one medium confidence enzyme (Ornithine carbamoyltransferase *argI*) that was a lower coverage match (78%) than the high confidence threshold (>80%). There is more evidence for a potential breakdown of essential amino acid synthesis in *P. hayleyella. P. hayleyella* had four potential gaps in its amino acid biosynthesis pathways. The L-isoleucine, L-leucine and L-valine pathways shared a single medium confidence enzyme candidate that is potentially a L-arabonate dehydratase rather than the necessary dihydroxy-acid dehydratase *ilvD* based on ublast bit scores. The L-tryptophan pathway had two medium confidence enzyme candidates for phosphoribosylanthranilate isomerase (*PRAI*), and the better scoring one was a lower coverage match (71%) than the high confidence threshold. However, given the degree of genome reduction that has already occurred in the reduced genome *D. discoideum*-symbionts, we consider it unlikely that the symbiotic relationship is based on amino acid exchange as essential amino acid synthesis pathways appear largely intact.

### D. discoideum-symbiont genomes share few horizontally transferred genetic elements

We looked for evidence of shared horizontally transmitted genetic elements. We identified 38, 29, and 27 genomic islands in each *D. discoideum*-symbiont genome (*P. agricolaris, P. bonniea*, and *P. hayleyella*), but none of the predicted genomic islands were closely related to a genomic island in another *D. discoideum*-symbiont genome. We found 133, 109, and 120 individually horizontally transferred genes in each *D. discoideum*-symbiont genome. One candidate was shared among all three genomes (type VI secretion system contractile sheath, large subunit) while two additional candidates were shared by *P. bonniea* and P. *hayleyella* (PIN family putative toxin-antitoxin system, toxin component; class I SAM-dependent methyltransferase). The scarcity of easily-identified shared horizontally transferred genetic elements suggest it is unlikely that a recent horizontal gene transfer event substantially contributed to the shared ability of these symbionts to persistently infect *D. discoideum*. If such an event had occurred, any such genes seem to have experienced amelioration over evolutionary time and cannot easily be distinguished from the rest of the genome (Lawrence & Ochman 1997).

### Shared secretion systems may mediate D. discoideum-Paraburkholderia symbiont interactions

Bacterial secretion systems are frequently implicated in host-symbiont interactions (Tseng et al. 2009; Coombes 2009). All *D. discoideum*-symbiont genomes possessed multiple type III secretions systems (T3SS) and type VI secretion systems (T6SS) in larger numbers than several of the other *Paraburkholderia* genomes examined (Figure 5; Table S4). Classification of T3SS showed that one specific T3SS operon shared among *D. discoideum*-symbionts falls into category 8 T3SS (Figure 6 & S4). This category of T3SS also includes BurBor found in the plant pathogen *Robbsia* (previously *Burkholderia*) *andropogonis* (Mannaa et al. 2019), as well as *Bordatella* species that include mammalian pathogens (Kamanova 2020). In addition, one specific T6SS operon is shared among *D. discoideum*-symbiont genomes and belongs to category i1 (Figure 7 & S5). More importantly, this T6SS operon clusters together with the virulence-causing T6SS-5 operon found in *Burkholderia mallei, B. pseudomallei*, and *B. thailandensis* (Lennings et al. 2019). *B. mallei* causes glanders disease and is an obligate pathogen that evolved from an ancestor shared with melioidosis-causing soil bacterium *B. pseudomallei* (Schell et al. 2007; Burtnick et al. 2011; Losada et al. 2010). *B. thailandensis* is sister species to the other two, and is a facultative pathogen similar to *B. pseudomallei* but with much lower clinical virulence (Lennings et al. 2019).

**Figure 5.**
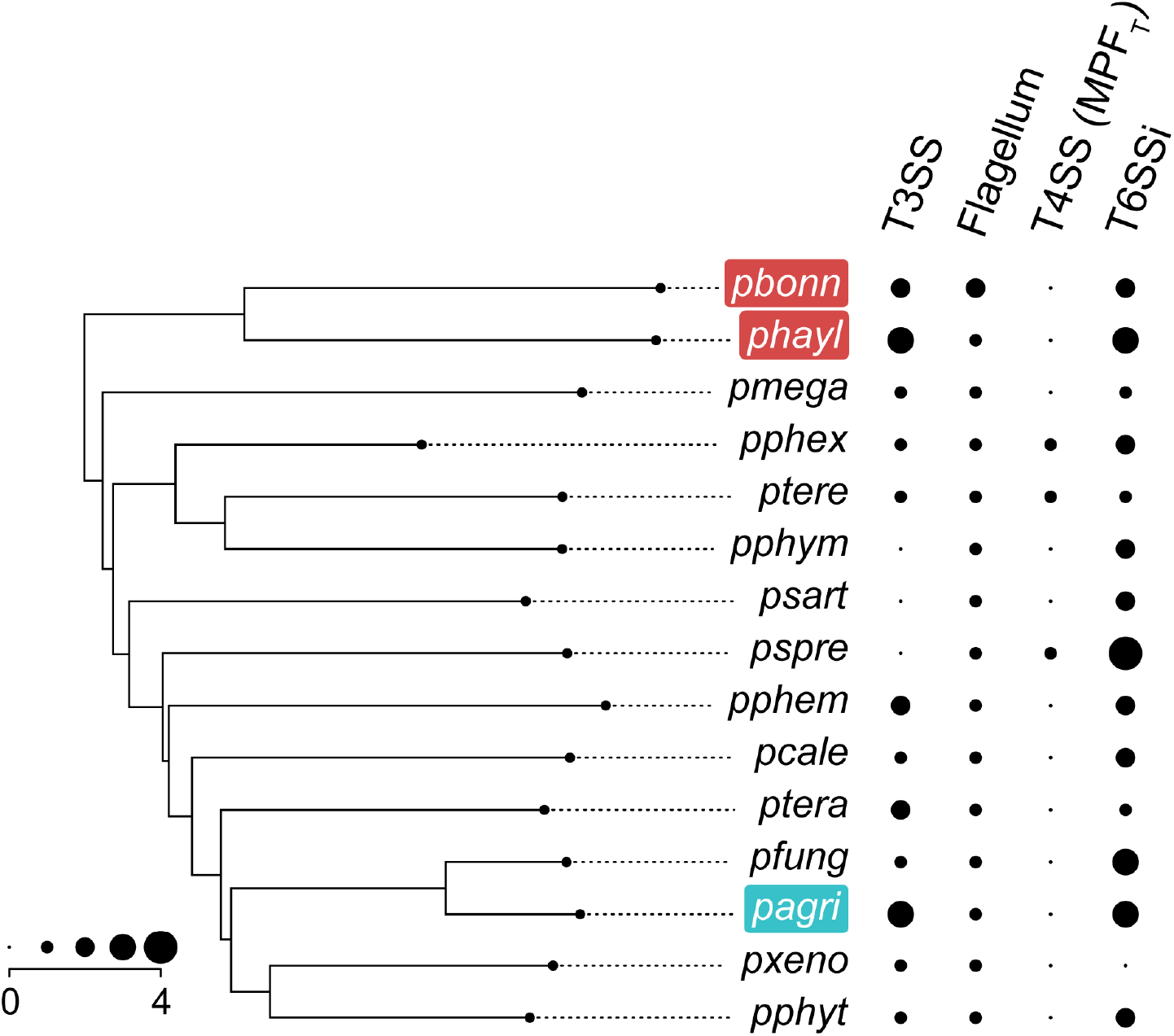
The abundances of secretion systems detected in *D. discoideum*-symbiont genomes and other *Paraburkholderia*. For the Type 4 Secretion System, only protein secretion (as opposed to conjugation-related) T4SS abundances are shown. The phylogeny is a species tree based on Brock et al 2020. (pagri = *P. agricolaris*; pbonn = *P. bonniea*; phayl = *P. hayleyella*; pcale = *P. caledonica*; pfung = *P. fungorum*, pmega = *P. megapolitana*, pphem = *P. phenazinium*; pphex = *P. phenoliruptrix*; pphym = *P. phymatum*; pphyt = *P. phytofirmans*; psart = *P. sartisoli*; pspre = *P. sprentiae*; ptera = *P. terricola*; ptere = *P. terrae*; pxeno = *P. xenovorans*).

**Figure 6.**
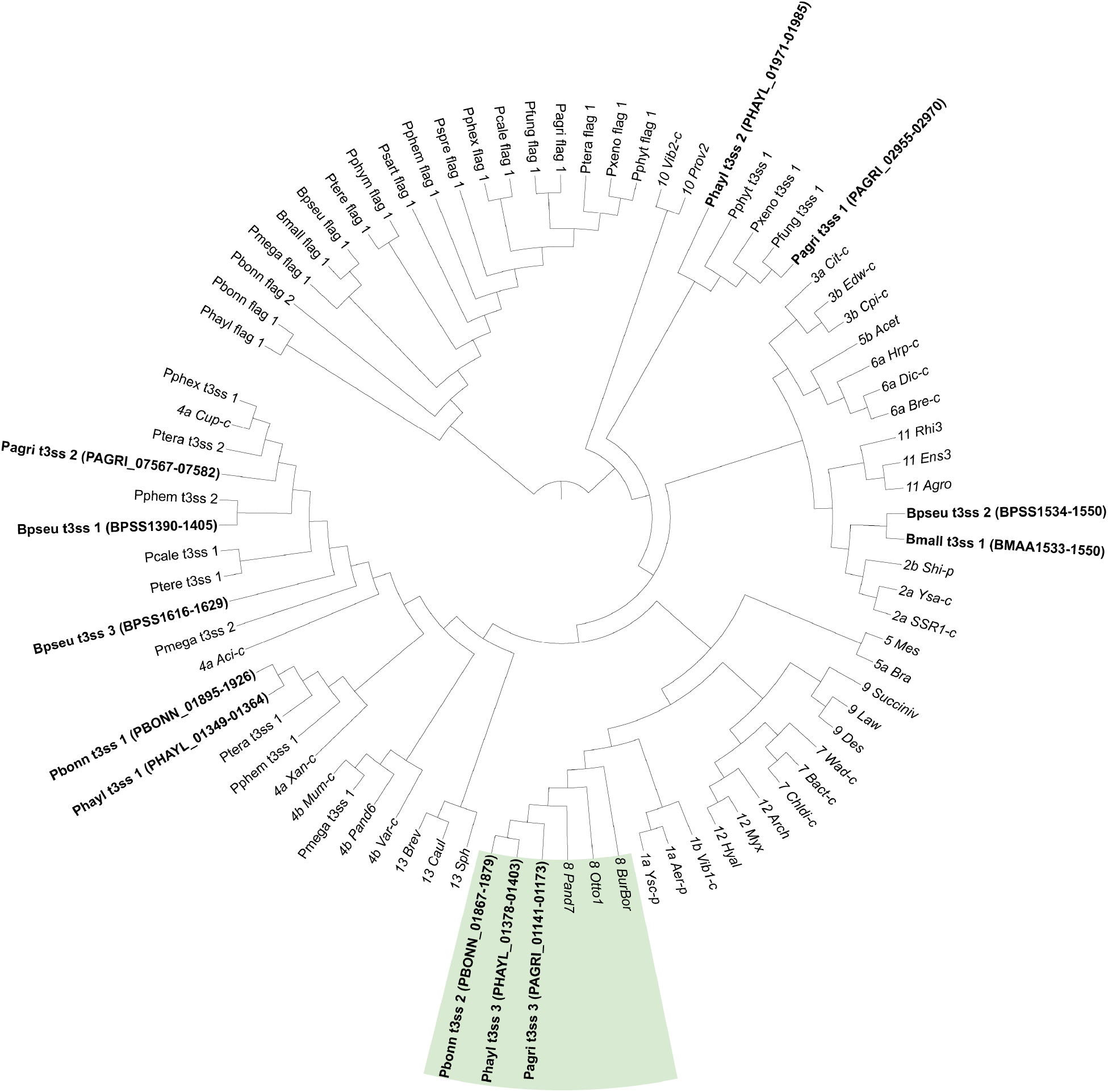
Type 3 secretion systems and flagella categorized using the conserved component genes sctJ (inner membrane ring; IPR003282), sctN (ATPase; IPR005714), and sctV (export apparatus; IPR006302). Branch lengths were ignored to improve readability of the ASTRAL tree topology. T3SS categories precede the name of the operon (e.g. “8 Pand7” is operon Pand7 belonging to category 8), downloaded from T3Enc database v1.0 (Hu et al. 2017). Tip labels for T3SS in the three *D. discoideum*-symbiont genomes and in *B. mallei* and *B. pseudomallei* are shown in bold font face with gene IDs for ease of reference. The clade containing the shared T3SS operon is shaded.

**Figure 7.**
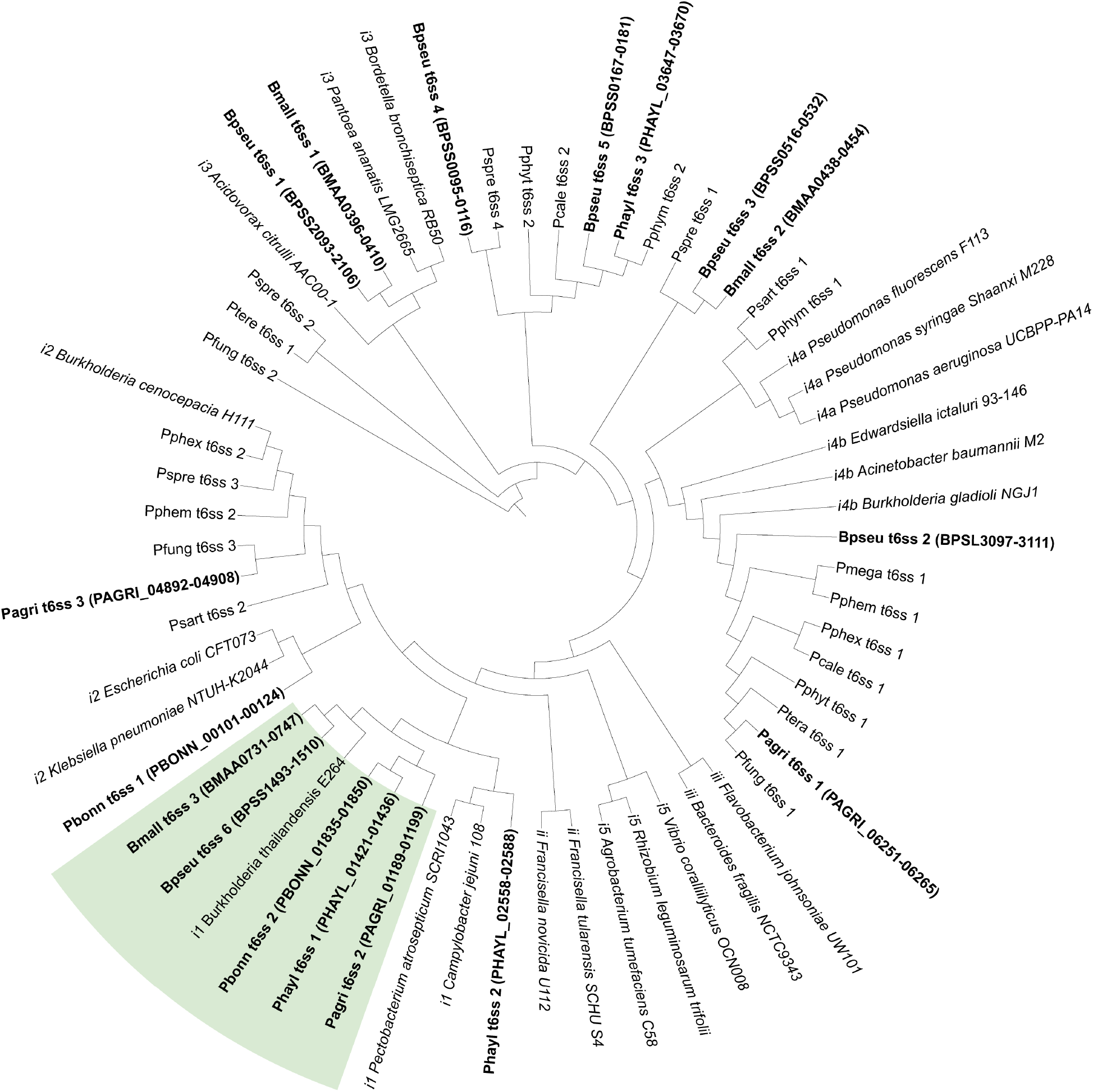
Type 6 secretion systems categorized using the conserved component genes tssB (sheath; COG3516), tssC (sheath; COG3517), and tssF (baseplate; COG3519). Branch lengths were ignored to improve readability of the ASTRAL tree topology. T6SS categories precede the name of the strain to which the operon belongs (e.g. “ii Francisella novicida U112” is belongs to category ii), downloaded from SecReT6 database v3.0 (Li et al. 2015). T6SS in the three *D. discoideum*-symbiont genomes and in *B. mallei* and *B. pseudomallei* are shown in bold font face with gene IDs for ease of reference. The clade containing the shared T6SS operon is shaded.

The T6SS-5-like and BurBor-like T3SS operons shared by the *D. discoideum*-symbionts are found directly next to each other on the respective genomes of *P. agricolaris, P. bonniea*, and *P. hayleyella. B. pseudomallei* and *B. thailandensis* also have a T3SS (T3SS-3 in the literature) adjacent to their T6SS-5 but these T3SS-3 operons appear to be unrelated to the BurBor-like T3SS found in the *D. discoideum*-symbiont genomes. It is worth noting that the two adjacent T3SS-3 and T6SS-5 operons in *B. pseudomallei* and *B. thailandensis* have been shown to be functionally linked and necessary for virulence, with the T3SS-3 effectors regulating the expression of the adjacent T6SS-5 (Sun et al. 2010; Chen et al. 2011; Schwarz et al. 2010; French et al. 2011).

We attempted to identify effector proteins that might be functionally linked to these *D. discoideum*-symbiont secretion systems (Table S5). We identified homologs of the T6SS effector VgrG-5 that would likely be associated with the shared T6SS-5-like operon (Table S5). Unexpectedly, VgrG-5 in *P. agricolaris* (gene ID PAGRI_01155) is a homolog but not an ortholog to VgrG-5 in the two reduced genomes (PBONN_01842 and PHAYL_01429). This suggests the possibility of two independent evolutionary origins of this T6SS effector, and potentially different functional roles. In *Burkholderia thailandensis*, VgrG-5 is necessary for post-infection cell-to-cell spread within mammalian hosts (Schwarz et al. 2014). For each genome, we predicted additional secretion system effectors, including a chaperonin ClpB and a sodium/solute symporter for *P. agricolaris*, and RHS (rearrangement hotspot) proteins that may mediate contact-dependent growth inhibition during bacterial competition for *P. hayleyella*. Lastly, we predicted secreted effectors containing eukaryotic domains specific to our Pfam clans of interest (Ank, TPR, LRR, Pentapeptide, F-box, and RING). Previous investigations of amoeba symbiont genomes have observed enrichment of proteins possessing these domains that hypothetically mediate physiological interactions with a eukaryotic host (Schmitz-Esser et al. 2010; Gomez-Valero & Buchrieser 2019; Schulz et al. 2016). Notably, two proteins each directly adjacent to VgrG-5 in *P. agricolaris* (PAGRI_01156-7) and *P. bonniea* (PBONN_01840-1) each contained pentapeptide repeat domains. InterProScan searches indicated that two proteins in *P. hayleyella* (PHAYL_01430-1) adjacent to VgrG-5 also contain pentapeptide repeat domains. However, no known functions are predicted for these protein pairs.

## Conclusion

The genomes of *Paraburkholderia* symbionts of *D. discoideum* present a unique opportunity to compare the significantly differently-sized genomes of three symbiont species that share the ability to persistently infect *D. discoideum*. We find evidence that relative to the other *Paraburkholderia* genomes we investigated, all three *D. discoideum*-symbiont genomes have increased secretion system and motility genes that potentially mediate interactions with their host. Specifically, adjacent type 3 and type 6 secretion system operons shared across all three *D. discoideum*-symbiont genomes may have an important role. The BurBor-like T3SS operon is closely related to one found in the plant pathogen *Robbsia andropogonis*. It includes a needle apparatus uncommon among *Burkholderia* T3SS that is used to inject rhizobitoxine into a wide range of plant hosts (Wallner et al. 2021; Mannaa et al. 2019). The adjacent T6SS operon is closely related to T6SS-5 shared by *B. mallei, B. pseudomallei*, and *B. thailandensis*. T6SS-5 is functionally important for the intercellular lifecycle of these pathogenic *Burkholderia* (Schwarz et al. 2014). We hypothesize that the BurBor-like T3SS operon is used during initial host infection and the T6SS-5-like operon may have a functional role post-infection. We also find orthologs to the T6 effector VgrG-5 specific to T6SS-5, as well as two neighboring potential effectors with eukaryote-like pentapeptide repeat domains in the three *D. discoideum*-symbiont genomes. Some but not all of the component genes of the shared T6SS-5-like and BurBor-like T3SS operons are among the 120 *D. discoideum*-symbiont-shared genes not found in any of the other *Paraburkholderia* genomes we compared. It is intriguing to consider the possibility that these genes were transferred among symbiont genomes within the *D. discoideum* amoeba host environment. Different *D. discoideum*-symbiont species have been found coinfecting amoeba hosts (Haselkorn et al. 2019), and diverse *Acanthamoeba* symbionts appear to share genes with each other that are functionally enriched for host interaction (Wang & Wu 2017).

While the secretion system features shared among *Paraburkholderia* symbionts of *D. discoideum* are striking, *P. agricolaris* is otherwise difficult to distinguish from other *Paraburkholderia* based on its genome size and content. However, the two reduced genomes of *P. bonniea* and *P. hayleyella* display characteristics that support their evolution in a host environment. All three species retain the ability to live outside of *D. discoideum*, but the genomes of *P. bonniea* and *P. hayleyella* show fewer transcriptional regulators, as well as fewer carbohydrate and inorganic ion transporters. The reduced genomes possess a combination of genes with molecular evolution patterns that indicate specific responses to the host environment (both stronger and weaker evolutionary constraints) rather than uniform deterioration under genetic drift. In addition, the lack of IS element proliferation and absence of excessive pseudogene accumulation compared to other *Paraburkholderia* genomes indicate that these already reduced genomes are relatively stable.

These combined pieces of evidence supports that the reduced genome *D. discoideum*-symbionts are professional symbionts specifically adapted to their protists hosts. We adopt the term “professional symbiont” from Husnik and colleages (2021) to refer to symbiont lineages that are ancestrally adapted to their specific hosts, that possess compact and streamlined genomes. Accordingly, we hypothesize that the symbiotic relationship between *D. discoideum* and the species with reduced genomes is persistent and potentially quite old. However, given the short generation time of protists it is entirely possible that what we call “old” is not as ancient as the symbioses and similarly stable stages of genome reduction observed in microbial symbionts of multicellular eukaryotes. In contrast to *P. bonniea* or *P. hayleyella*, intraspecific genetic variation appears to be larger for *P. agricolaris* (Haselkorn et al. 2019), suggesting that *P. agricolaris* host adaptation may be ongoing and more dynamic. We look forward to expanding these analyses to a larger collection of *D. discoideum*-symbiont genomes in the future, to identify both convergent and divergent host adaptation patterns among *D. discoideum*-symbionts and to continue to add to a growing body of work across diverse protist-prokaryote symbioses.

## Acknowledgements

We thank students and colleagues from the Noh lab (Anna Chen, Kayla Dixon, Laura Drepanos) and Strassmann-Queller lab (Tammy Haselkorn, Clarissa Dzikunu) who contributed to aspects of this project during its development. This work is supported by the National Science Foundation under Grant Numbers IOS 1656756 and DEB 1753743 (JS and DQ), and by the National Institutes of Health and its National Institute of General Medical Sciences by an Institutional Development Award (IDeA) under Grant Numbers P20GM103423 (subaward to SN) and Colby College startup funds (SN).

## Notes

https://github.com/noh-lab/comparative-dicty-symbionts

### Competing Interest Statement

The authors have declared no competing interest.

