## Supplemental Tables for "Comparative genomics support reduced-genome *Paraburkholderia* symbionts of *Dictyostelium discoideum* amoebas are ancestrally adapted professional symbionts"

Supplementary Table 1. *Paraburkholderia* genomes examined in this study

| Genome (species and strain) | Abbreviation | Category | RefSeq assembly accession |
| --- | --- | --- | --- |
| *P. agricolaris* BaQS159 | PAGRI | *D. discoideum*-symbiont | GCF_009455635.1_ASM945563v1_genomic.fna |
| *P. bonniea* BbQS859 | PBONN | *D. discoideum*-symbiont | GCF_009455625.1_ASM945562v1_genomic.fna |
| *P. hayleyella* BhQS11 | PHAYL | *D. discoideum*-symbiont | GCF_009455685.1_ASM945568v1_genomic.fna |
| *P. fungorum* ATCC BAA-463 | PFUNG | Symbiotic | GCF_000961515.1_ASM96151v1_genomic.fna |
| *P. megapolitana* LMG23650 | PMEGA | Symbiotic | GCF_007556815.1_ASM755681v1_genomic.fna |
| *P. phenoliruptrix* BR3459a | PPHEX | Symbiotic | GCF_000300095.1_ASM30009v1_genomic.fna |
| *P. phymatum* STM815 | PPHYM | Symbiotic | GCF_000020045.1_ASM2004v1_genomic.fna |
| *P. phytofirmans* PsJN | PPHYT | Symbiotic | GCF_000020125.1_ASM2012v1_genomic.fna |
| *P. sprentiae* WSM5005 | PSPRE | Symbiotic | GCF_001865575.1_ASM186557v1_genomic.fna |
| *P. caledonica* PHRS4, | PCALE | Free-living | GCF_003330745.1_ASM333074v1_genomic.fna |
| *P. phenazinium* LMG2247 | PPHEM | Free-living | GCF_900100735.1_IMG2651870170_genomic.fna |
| *P. sartisoli* LMG24000 | PSART | Free-living | GCF_900107685.1_IMG2651870102_genomic.fna |
| *P. terricola* mHS1 | PTERA | Free-living | GCF_003330825.1_ASM333082v1_genomic.fna |
| *P. terrae* DSM17804 | PTERE | Free-living | GCF_002902925.1_ASM290292v1_genomic.fna |
| *P. xenovorans* LB400 | PXENO | Free-living | GCF_000756045.1_ASM75604v1_genomic.fna |

Supplementary Table 2. Insertion Sequence (IS) elements found in *D. discoideum*-symbiont *Paraburkholderia* genomes

| Genome | IS1090  (IS256 family) | ISBmu21  (IS6 family) | ISBp1  (IS3 family) | ISBuph1  (IS5 family) | ISPa37  (IS30 family) |
| --- | --- | --- | --- | --- | --- |
| *P. agricolaris* BaQS159 | 6 | 1 | 0 | 0 | 0 |
| *P. bonniea* BbQS859 | 0 | 0 | 2 | 3 | 0 |
| *P. hayleyella* BhQS11 | 0 | 0 | 0 | 0 | 1 |

Supplementary Table 3. Best scoring candidates for predicted amino acid biosynthesis pathways in *D. discoideum* -symbiont *Paraburkholderia* genomes

| Amino acid | Gene | *P. agricolaris BaQS159* | *P. bonniea BbQS859* | *P. hayleyella BhQS11* | Description |
| --- | --- | --- | --- | --- | --- |
| arg | *argA* | PAGRI_01412 | PBONN_01141 | PHAYL_02229 | N-acylglutamate synthase |
|  | *argB* | PAGRI_00116 | PBONN_00046 | PHAYL_00082 | N-acylglutamate kinase |
|  | *argC* | PAGRI_04578 | PBONN_00437 | PHAYL_00446 | N-acylglutamylphosphate reductase |
|  | *argD* | PAGRI_02772 | PBONN_01729 | PHAYL_01489 | N-acetylornithine aminotransferase |
|  | *argE* | PAGRI_01357 | PBONN_01537 | PHAYL_01001 | N-acetylornithine deacetylase |
|  | *carA* | PAGRI_01014 | PBONN_02041 | PHAYL_02324 | carbamoyl phosphate synthase subunit alpha |
|  | *carB* | PAGRI_01015 | PBONN_02040 | PHAYL_02323 | carbamoyl phosphate synthase subunit beta |
|  | *argI* | PAGRI_00606 | PBONN_02351* | PHAYL_02432 | ornithine carbamoyltransferase |
|  | *argG* | PAGRI_00605 | PBONN_02353 | PHAYL_02433 | arginosuccinate synthetase |
|  | *argH* | PAGRI_03246 | PBONN_01037 | PHAYL_01059 | arginosuccinate lyase |
| gln | *gltX* | PAGRI_01057 | PBONN_01058 | PHAYL_01095 | glutamyl-tRNA(Glx) synthetase |
|  | *glnA* | PAGRI_01420 | PBONN_01149 | PHAYL_02222 | glutamine synthetase |
|  | *gatA* | PAGRI_00106 | PBONN_00037 | PHAYL_00072 | glutamyl-tRNA(Gln) amidotransferase subunit A |
|  | *gatB* | PAGRI_00107 | PBONN_00038 | PHAYL_00073 | glutamyl-tRNA(Gln) amidotransferase subunit B |
|  | *gatC* | PAGRI_00105 | PBONN_00036 | PHAYL_00071 | glutamyl-tRNA(Gln) amidotransferase subunit C |
| pro | *proB* | PAGRI_03791 | PBONN_02585 | PHAYL_02797 | glutamate 5-kinase |
|  | *proA* | PAGRI_00426 | PBONN_02536 | PHAYL_02750 | gamma-glutamylphosphate reductase |
|  | *proC* | PAGRI_03641 | PBONN_02465 | PHAYL_02667 | pyrroline-5-carboxylate reductase |
| chorismate | *aroG* | PAGRI_03631 | PBONN_02456 | PHAYL_02659 | 3-deoxy-7-phosphoheptulonate synthase |
|  | *aroB* | PAGRI_03945 | PBONN_00516 | PHAYL_00530 | 3-dehydroquinate synthase |
|  | *aroD* | PAGRI_03770 | PBONN_02566 | PHAYL_02778 | 3-dehydroquinate dehydratase |
|  | *aroE* | PAGRI_05515 | PBONN_02560 | PHAYL_03196 | shikimate dehydrogenase |
|  | *aroL* | PAGRI_03946 | PBONN_00515 | PHAYL_00529 | shikimate kinase |
|  | *aroA* | PAGRI_00874 | PBONN_00985 | PHAYL_01015 | 3-phosphoshikimate 1-carboxyvinyltransferase |
|  | *aroC* | PAGRI_01111 | PBONN_01353 | PHAYL_02013 | chorismate synthase |
| phe/tyr | *cmutase* | PAGRI_00871 | PBONN_00982 | PHAYL_01012 | chorismate mutase |
| phe | *pre-dehydratase* | PAGRI_00871 | PBONN_00982 | PHAYL_01012 | prephenate dehydratase |
|  | *ilvE* | PAGRI_00872 | PBONN_00983 | PHAYL_01013 | aromatic amino acid aminotransferase |
| tyr | *pre-dehydratase* | PAGRI_00873 | PBONN_00984 | PHAYL_01014 | prephenate dehydrogenase |
|  | *tyrB* | PAGRI_00872 | PBONN_00983 | PHAYL_01013 | aromatic amino acid aminotransferase |
| trp | *trpE* | PAGRI_03847 | PBONN_00576 | PHAYL_00589 | anthranilate synthase subunit TrpE |
|  | *trpD_1* | PAGRI_03848 | PBONN_00575 | PHAYL_00586 | glutamine amidotransferase of anthranilate synthase |
|  | *trpD_2* | PAGRI_03849 | PBONN_00574 | PHAYL_00585 | anthranilate phosphoribosyltransferase |
|  | *PRAI* | PAGRI_07665 | PBONN_03508 | PHAYL_03576* | phosphoribosylanthranilate isomerase |
|  | *IGPS* | PAGRI_03850 | PBONN_00573 | PHAYL_00584 | indole-3-glycerol phosphate synthase |
|  | *trpA* | PAGRI_07662 | PBONN_03505 | PHAYL_03573 | indoleglycerol phosphate aldolase |
|  | *trpB* | PAGRI_07664 | PBONN_03507 | PHAYL_03575 | tryptophan synthase |
| asn | *aspS2* | PAGRI_03743 | PBONN_00648 | PHAYL_00666 | aspartyl-tRNA(Asp/Asn) synthetase |
|  | *gatA* | PAGRI_00106 | PBONN_00037 | PHAYL_00072 | glutamyl-tRNA(Gln) amidotransferase subunit A |
|  | *gatB* | PAGRI_00107 | PBONN_00038 | PHAYL_00073 | glutamyl-tRNA(Gln) amidotransferase subunit B |
|  | *gatC* | PAGRI_00105 | PBONN_00036 | PHAYL_00071 | glutamyl-tRNA(Gln) amidotransferase subunit C |
| lys | *asp_kinase* | PAGRI_01585 | PBONN_01273 | PHAYL_02093 | aspartate kinase |
|  | *asd* | PAGRI_07668 | PBONN_03511 | PHAYL_03580 | asparate semi-aldehyde dehydrogenase |
|  | *dapA* | PAGRI_07296 | PBONN_01227 | PHAYL_02141 | 4-hydroxy-tetrahydrodipicolinate synthase |
|  | *dapB* | PAGRI_00420 | PBONN_02544 | PHAYL_02756 | 4-hydroxy-tetrahydrodipicolinate reductase |
|  | *dapD* | PAGRI_01631 | PBONN_01775 | PHAYL_01541 | tetrahydrodipicolinate succinylase |
|  | *dapC* | PAGRI_01632 | PBONN_01774 | PHAYL_01542 | N-succinyldiaminopimelate aminotransferase |
|  | *dapE* | PAGRI_01629 | PBONN_01777 | PHAYL_01539 | succinyl-diaminopimelate desuccinylase |
|  | *dapF* | PAGRI_00127 | PBONN_00057 | PHAYL_00093 | diaminopimelate epimerase |
|  | *lysA* | PAGRI_04401 | PBONN_00506 | PHAYL_00522 | diaminopimelate decarboxylase |
| met/thr | *asp-kinase* | PAGRI_01585 | PBONN_01273 | PHAYL_02093 | aspartate kinase |
|  | *asd* | PAGRI_07668 | PBONN_03511 | PHAYL_03580 | aspartate semi-aldehyde dehydrogenase |
|  | *hom* | PAGRI_02315 | PBONN_01485 | PHAYL_01903 | homoserine dehydrogenase |
|  | *metA* | PAGRI_00113 | PBONN_00042 | PHAYL_00078 | homoserine O-succinyltransferase |
| met | *metZ* | PAGRI_07656 | PBONN_03498 | PHAYL_03567 | O-succinylhomoserine sulfhydrylase |
|  | *metH* | PAGRI_00253, PAGRI_00254 | PBONN_02806, PBONN_02805 | PHAYL_02945, PHAYL_02944 | vitamin B12-dependent methionine synthase |
|  | *B12_reactivation_domain* | PAGRI_00253 | PBONN_02806 | PHAYL_02945 | MetH reactivation domain |
| thr | *thrB* | PAGRI_07616 | PBONN_01954 | PHAYL_01321 | homoserine kinase |
|  | *thrC* | PAGRI_02314 | PBONN_01486 | PHAYL_01902 | threonine synthase |
| ile | *ilvA* | PAGRI_03754 | PBONN_00637 | PHAYL_00654 | threonine deaminase |
| ile/leu/val | *ilvH* | PAGRI_03065 | PBONN_03362 | PHAYL_03450 | acetohydroxybutanoate synthase catalytic subunit |
|  | *ilvI* | PAGRI_03064 | PBONN_03361 | PHAYL_03449 | acetohydroxybutanoate synthase regulatory subunit |
|  | *ilvC* | PAGRI_03063 | PBONN_03360 | PHAYL_03448 | 2-hydroxy-3-ketol-acid reductoisomerase |
|  | *ilvD* | PAGRI_03332 | PBONN_02205 | PHAYL_03298* | (R)-2,3-dihydroxy-3-methylpentanoate dehydratase |
|  | *ilvE* | PAGRI_00529 | PBONN_02403 | PHAYL_02490 | aromatic amino acid aminotransferase |
| leu | *leuA* | PAGRI_01249 | PBONN_02670 | PHAYL_02850 | 2-isopropylmalate synthase |
|  | *leuC* | PAGRI_07672 | PBONN_03514 | PHAYL_03584 | isopropylmalate isomerase large subunit |
|  | *leuD* | PAGRI_07670 | PBONN_03513 | PHAYL_03582 | isopropylmalate isomerase small subunit |
|  | *leuB* | PAGRI_07669 | PBONN_03512 | PHAYL_03581 | 3-isopropylmalate dehydrogenase |
| cys | *cysE* | PAGRI_05168 | PBONN_02934 | PHAYL_03076 | serine acetyltransferase |
|  | *cysK* | PAGRI_00884 | PBONN_00995 | PHAYL_01029 | O-acetylserine sulfhydrylase |
| gly | *glyA* | PAGRI_03574 | PBONN_00808 | PHAYL_00862 | serine hydroxymethyltransferase |
| ser | *serA* | PAGRI_03753 | PBONN_00638 | PHAYL_00656 | 3-phosphoglycerate dehydrogenase |
|  | *serC* | PAGRI_00870 | PBONN_00981 | PHAYL_01011 | 3-phosphoserine aminotransferase |
|  | *serB* | PAGRI_02265 | PBONN_01530 | PHAYL_01843 | phosphoserine phosphatase |
| his | *prs* | PAGRI_00358 | PBONN_02660 | PHAYL_02840 | ribose-phosphate diphosphokinase |
|  | *hisG* | PAGRI_03910 | PBONN_00545 | PHAYL_00556 | ATP phosphoribosyltransferase |
|  | *hisI* | PAGRI_03901 | PBONN_00553 | PHAYL_00564 | phosphoribosyl-ATP pyrophosphatase |
|  | *hisE* | PAGRI_03902 | PBONN_00552 | PHAYL_00563 | phosphoribosyl-AMP cyclohydrolase |
|  | *hisA* | PAGRI_03904 | PBONN_00550 | PHAYL_00561 | isomerase HisA |
|  | *hisF* | PAGRI_03903 | PBONN_00551 | PHAYL_00562 | IGP synthase |
|  | *hisH* | PAGRI_03905 | PBONN_00549 | PHAYL_00560 | IGP synthase |
|  | *hisB* | PAGRI_03907 | PBONN_00548 | PHAYL_00559 | IGP dehydratase |
|  | *hisC* | PAGRI_00355 | PBONN_00547 | PHAYL_00558 | histidinol-phosphate aminotransferase |
|  | *hisN* | PAGRI_03612 | PBONN_02434 | PHAYL_02637 | histidinol-phosphate phosphatase |
|  | *hisD* | PAGRI_03909 | PBONN_00546 | PHAYL_00557 | histidinol dehydrogenase |

* Medium confidence enzymes in GapMind analysis

Supplementary Table 4. Predicted secretion systems for each *Paraburkholderia* genome examined in this study

| genome | T1SS | T2SS | Tad | T3SS | Flagella | pT4SS | cT4SS | T5aSS | T5bSS | T5cSS | T6SSi |
| --- | --- | --- | --- | --- | --- | --- | --- | --- | --- | --- | --- |
| PAGRI | 0 | 1 | 1 | 3 | 1 | 0 | 1 | 2 | 2 | 3 | 3 |
| PBONN | 3 | 1 | 1 | 2 | 2 | 0 | 0 | 3 | 3 | 3 | 2 |
| PHAYL | 2 | 1 | 1 | 3 | 1 | 0 | 0 | 3 | 3 | 5 | 3 |
| PCALE | 2 | 1 | 0 | 1 | 1 | 0 | 0 | 0 | 2 | 3 | 2 |
| PFUNG | 1 | 1 | 2 | 1 | 1 | 0 | 0 | 2 | 5 | 2 | 3 |
| PMEGA | 4 | 1 | 1 | 1 | 1 | 0 | 0 | 0 | 6 | 3 | 1 |
| PPHEM | 3 | 1 | 2 | 2 | 1 | 0 | 0 | 1 | 1 | 5 | 2 |
| PPHEX | 2 | 1 | 2 | 1 | 1 | 1 | 2 | 0 | 7 | 4 | 2 |
| PPHYM | 2 | 1 | 1 | 0 | 1 | 0 | 1 | 0 | 5 | 5 | 2 |
| PPHYT | 1 | 1 | 2 | 1 | 1 | 0 | 3 | 1 | 2 | 3 | 2 |
| PSART | 2 | 1 | 1 | 0 | 1 | 0 | 1 | 3 | 3 | 1 | 2 |
| PSPRE | 2 | 1 | 0 | 0 | 1 | 1 | 1 | 1 | 6 | 3 | 4 |
| PTERA | 2 | 1 | 2 | 2 | 1 | 0 | 0 | 0 | 2 | 4 | 1 |
| PTERE | 2 | 1 | 1 | 1 | 1 | 1 | 0 | 0 | 5 | 3 | 1 |
| PXENO | 3 | 1 | 1 | 1 | 1 | 0 | 4 | 0 | 5 | 2 | 0 |

* T4SS are classified into protein secretion vs. conjugation-related types (pT4SS vs. cT4SS)

Supplementary Table 5. Secreted effectors predicted in *D. discoideum*-symbiont *Paraburkholderia* genomes

| Genome | Gene ID | Annotation | Annotation source |
| --- | --- | --- | --- |
| ***P. agricolaris* BaQS159** | PAGRI_00020 | Pentapeptide domain protein | effectiveELD |
|  | PAGRI_00362 | TPR domain protein | effectiveELD |
|  | PAGRI_00592 | TPR domain protein | effectiveELD |
|  | PAGRI_00623 | TPR domain protein | effectiveELD |
|  | PAGRI_01039 | Pentapeptide domain protein | effectiveELD |
|  | PAGRI_01112 | TPR domain protein | effectiveELD |
|  | PAGRI_01155 | T6SS effector VgrG-5 | VFDB |
|  | PAGRI_01156 | Pentapeptide domain protein | effectiveELD |
|  | PAGRI_01157 | Pentapeptide domain protein | effectiveELD |
|  | PAGRI_01179 | TPR domain protein | effectiveELD |
|  | PAGRI_01202 | LRR domain protein | effectiveELD |
|  | PAGRI_02308 | T6SS effector | BastionHub |
|  | PAGRI_02840 | TPR domain protein | effectiveELD |
|  | PAGRI_02920 | TPR domain protein | effectiveELD |
|  | PAGRI_03740 | TPR domain protein | effectiveELD |
|  | PAGRI_04081 | TPR domain protein | effectiveELD |
|  | PAGRI_04899 | T6SS effector VgrG-5 | VFDB |
|  | PAGRI_04916 | T4SS effector | BastionHub |
|  | PAGRI_05317 | T3SS effector | BastionHub |
|  | PAGRI_06262 | TPR domain protein | effectiveELD |
|  | PAGRI_06513 | TPR domain protein | effectiveELD |
|  | PAGRI_07504 | TPR domain protein | effectiveELD |
| ***P. bonniea* BbQS859** | PBONN_00263 | Ank domain protein | effectiveELD |
|  | PBONN_00282 | T1SS effector | BastionHub |
|  | PBONN_00789 | T3SS effector | BastionHub |
|  | PBONN_01051 | Pentapeptide domain protein | effectiveELD |
|  | PBONN_01823 | T1SS effector | BastionHub |
|  | PBONN_01840 | Pentapeptide domain protein | effectiveELD |
|  | PBONN_01841 | Pentapeptide domain protein | effectiveELD |
|  | PBONN_01842 | T6SS effector VgrG-5 | VFDB |
|  | PBONN_01861 | TPR domain protein | effectiveELD |
|  | PBONN_02315 | T1SS effector | BastionHub |
|  | PBONN_02901 | LRR domain protein | effectiveELD |
|  | PBONN_03205 | Pentapeptide domain protein | effectiveELD |
|  | PBONN_03247 | T6SS effector | BastionHub |
|  | PBONN_03375 | Ank domain protein | effectiveELD |
|  | PBONN_03418 | T1SS effector | BastionHub |
| ***P. hayleyella* BhQS11** | PHAYL_00296 | T1SS effector | BastionHub |
|  | PHAYL_00422 | T6SS effector | BastionHub |
|  | PHAYL_00425 | T6SS effector | BastionHub |
|  | PHAYL_00686 | T1SS effector | BastionHub |
|  | PHAYL_01212 | T6SS effector | BastionHub |
|  | PHAYL_01429 | T6SS effector VgrG-5 | VFDB |
|  | PHAYL_02569 | T6SS effector | BastionHub |
|  | PHAYL_02573 | T6SS effector | BastionHub |
|  | PHAYL_02589 | T6SS effector | BastionHub |
|  | PHAYL_02594 | T6SS effector | BastionHub |
|  | PHAYL_02596 | T6SS effector | BastionHub |
|  | PHAYL_02973 | T2SS effector | BastionHub |
|  | PHAYL_02975 | T6SS effector | BastionHub |
|  | PHAYL_02976 | T6SS_effector | BastionHub |
|  | PHAYL_03032 | Ank domain protein | effectiveELD |
