## Supplemental Figures for "Comparative genomics support reduced-genome *Paraburkholderia* symbionts of *Dictyostelium discoideum* amoebas are ancestrally adapted professional symbionts"

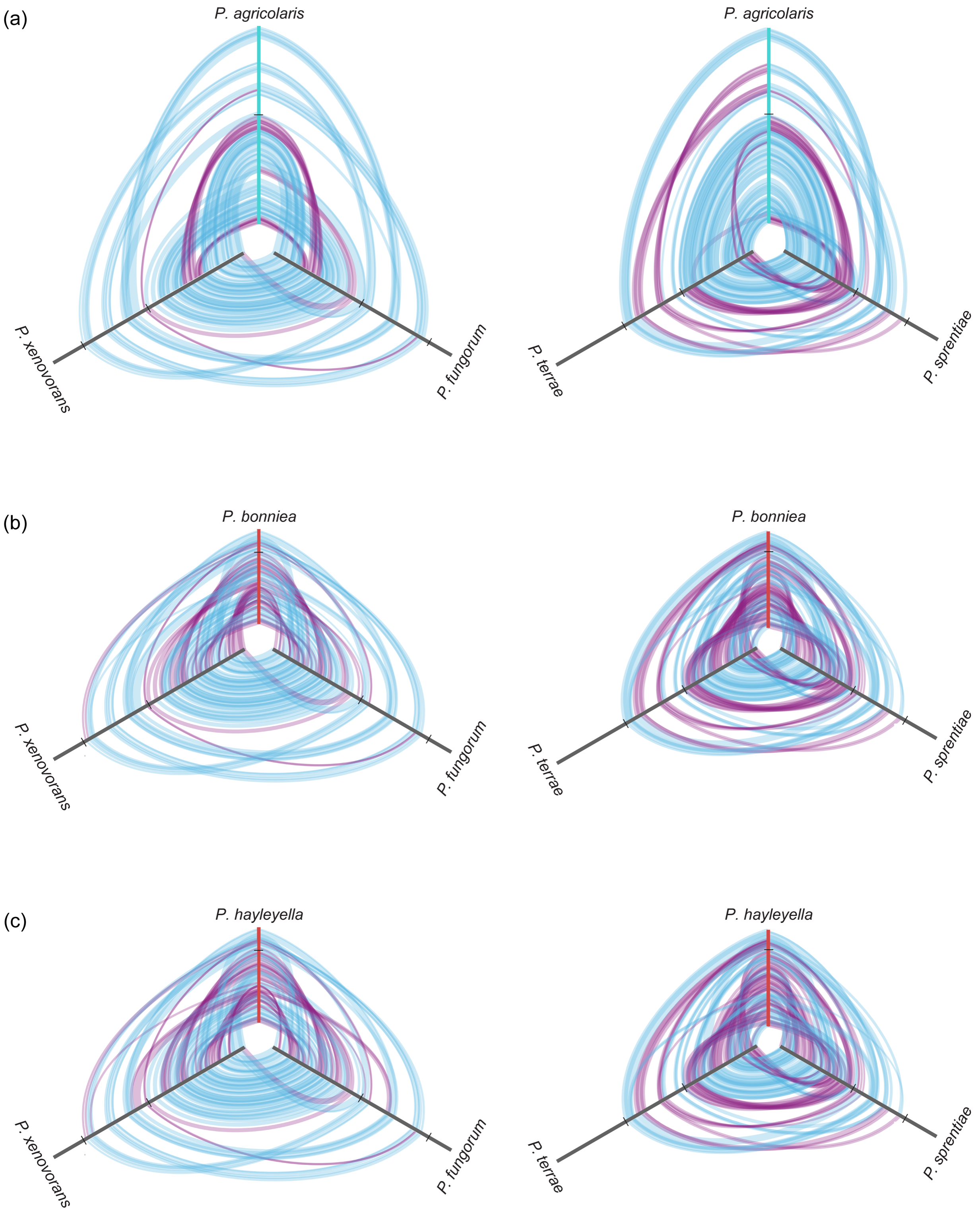


Figure S1. Hive plots of *D. discoideum*-symbiont genomes compared to four other *Paraburkholderia* genomes (a) *P. agricolaris*, (b) *P. bonniea*, and (c) *P. hayleyella*. Locally colinear blocks between pairs of genomes are shown as bands that connect the axes (genomes). Only blocks above the median size are shown for visual clarity. Alignment of locally colinear blocks are distinguished between forward (blue) and reverse (purple) orientation. Axes are oriented center out, and boundaries between chromosomes are shown as ticks. The two reduced genomes show considerable genome rearrangement while Chromosome 1 is largely similar for these *Paraburkholderia* genomes as indicated by the overall lack of gaps toward the center of each hive plot. The origin of chromosome 1 appears to be slightly different for other *Paraburkholderia* genomes (see connections between the center origin and regions close to the white tick mark indicating the boundary between linearized chromosones 1 and 2).

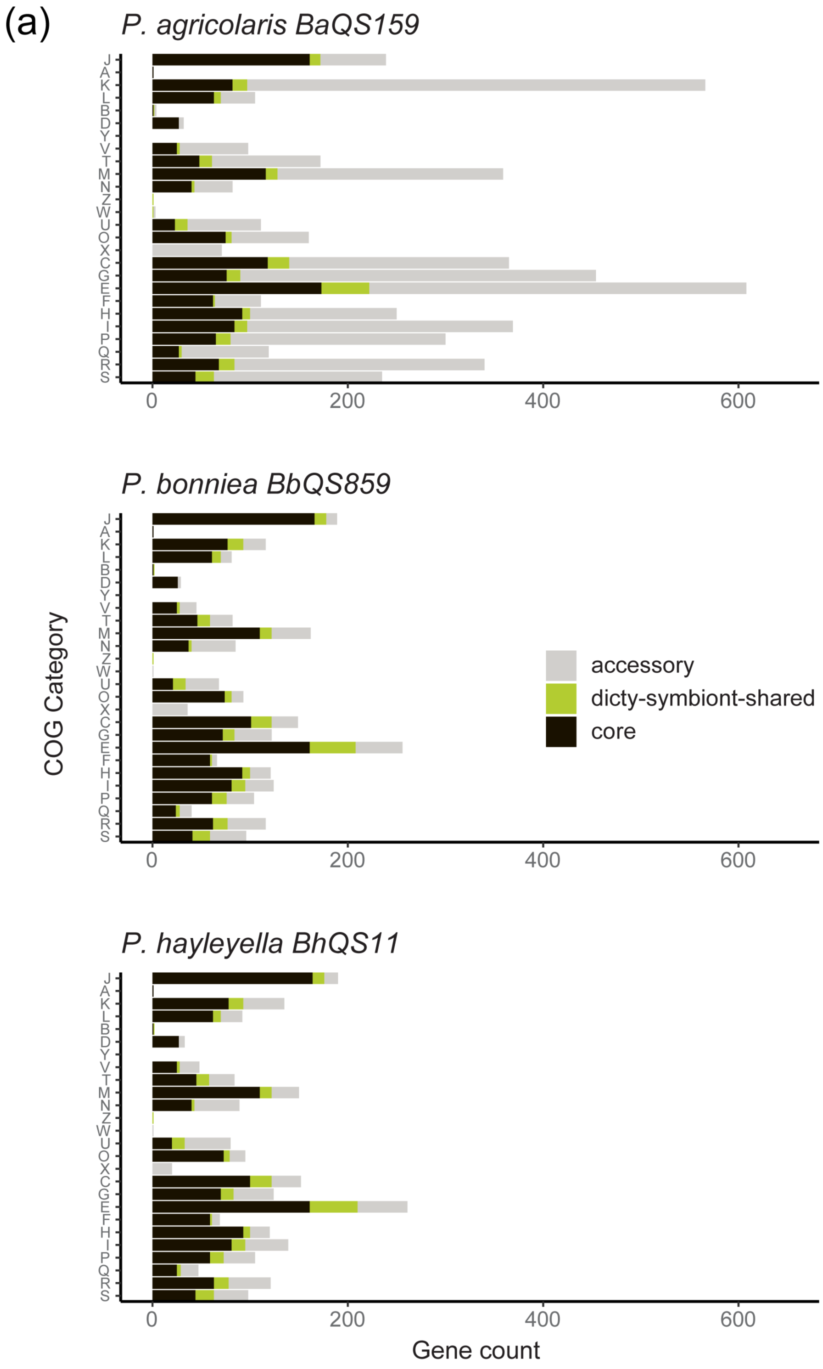

(a)
P. agricolaris BaQS159
P. bonniea BbQS859
P. hayleyella BhQS11
J
A
K
L
B
D
Y
V
T
M
N
Z
W
U
O
X
C
G
E
F
H
I
P
Q
R
S
0
200
400
600
COG Category
Gene count
accessory
dicty-symbiont-shared
core

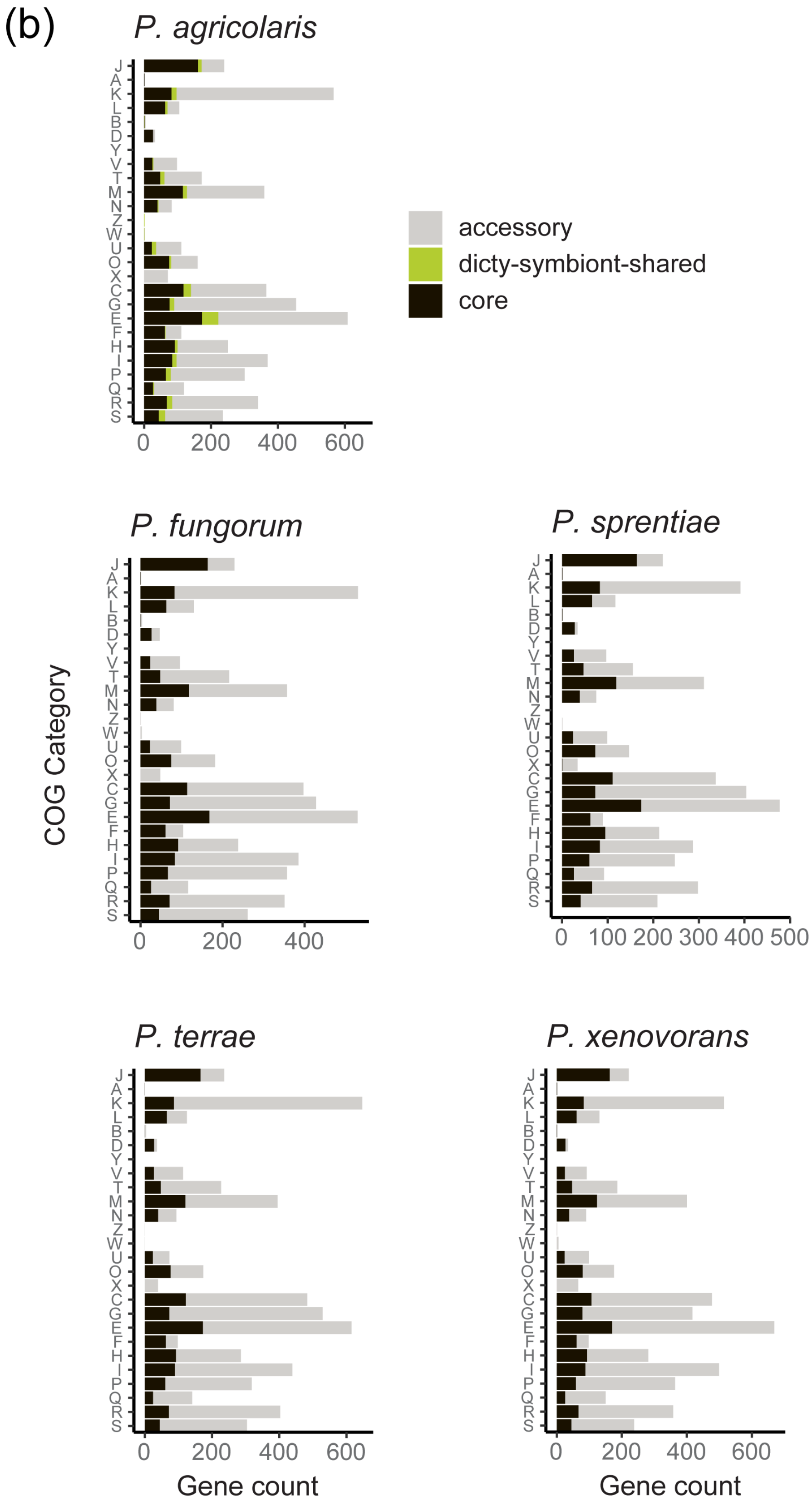

(b)
P. agricolaris
P. fungorum
P. sprentiae
P. terrae
P. xenovorans
accessory
dicty-symbiont-shared
core
COG Category
J
A
K
L
B
D
Y
V
T
M
N
Z
W
U
O
X
C
G
E
F
H
I
P
Q
R
S
0
200
400
600
100
300
500
Gene count

Figure S2. (a) Composition of each *D. discoideum*-symbiont genome divided into core genes shared among all *Paraburkholderia* examined, “dicty” genes shared among all three *D. discoideum*-symbiont genomes but excluding core genes, and all other accessory genes. (b) Compositions of *P. agricolaris* and 4 representative *Paraburkholderia* genomes divided into core genes shared among all *Paraburkholderia* examined, “dicty” genes shared among all three *D. discoideum*-symbiont genomes but excluding core genes, and all other accessory genes. (COG categories J = Translation, ribosomal structure and biogenesis; A = RNA processing and modification; K = Transcription; L = Replication, recombination and repair; B = Chromatin structure and dynamics; D = Cell cycle control, cell division, chromosome partitioning; Y = Nuclear structure; V = Defense mechanisms; T = Signal transduction mechanisms; M = Cell wall/ membrane/ envelope biogenesis; N = Cell motility; Z = Cytoskeleton; W = Extracellular structures; U = Intracellular trafficking, secretion, and vesicular transport O = Posttranslational modification, protein turnover, chaperones; X = Mobilome: prophages, transposons; C = Energy production and conversion; G = Carbohydrate transport and metabolism; E = Amino acid transport and metabolism; F = Nucleotide transport and metabolism; H = Coenzyme transport and metabolism; I = Lipid transport and metabolism; P = Inorganic ion transport and metabolism; Q = Secondary metabolites biosynthesis, transport and catabolism; R = General function prediction only; S = Function unknown)

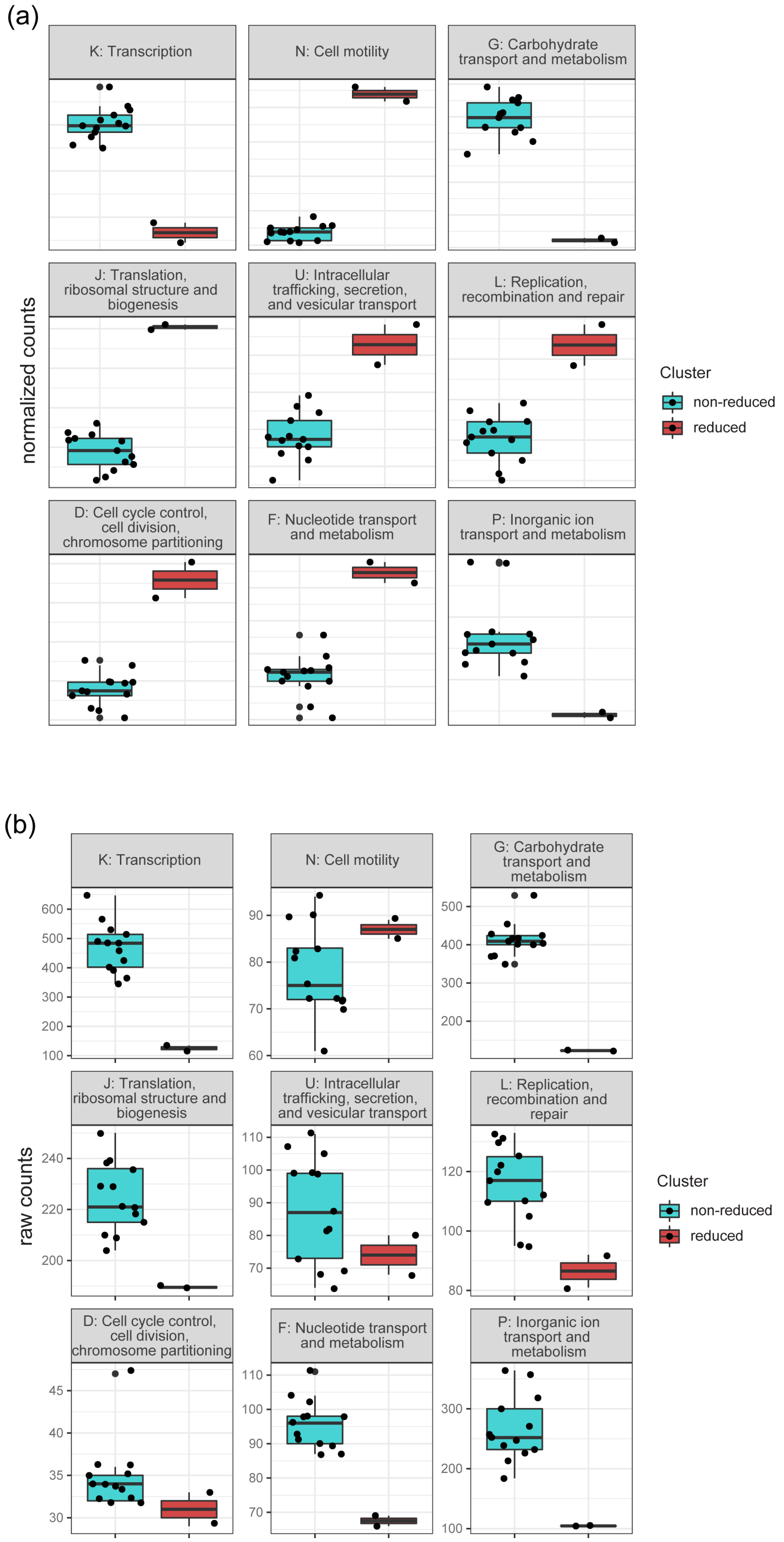


Figure S3. Comparison of significantly different COG functional categories between reduced and non-reduced Paraburkholderia genomes. Normalized (a) and raw (b) gene counts are shown. Categories K, N, G, and P were consistently different in the same direction for reduced genomes vs. non-reduced genomes.

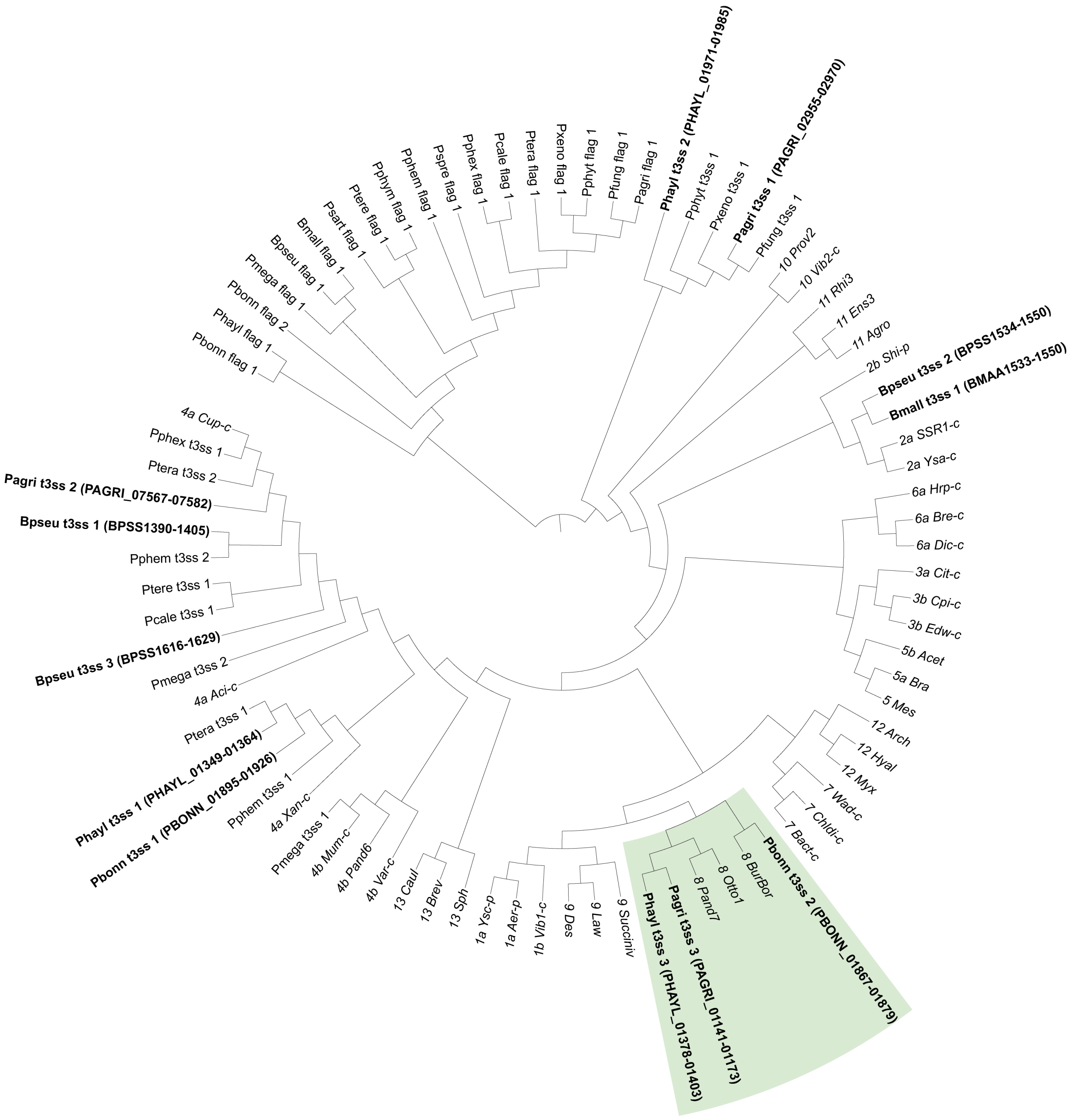


Figure S4. Type 3 secretion systems and flagella categorized using gene trees of the conserved component genes sctJ (inner membrane ring; IPR003282), sctN (ATPase; IPR005714), and sctV (export apparatus; IPR006302). Branch lengths were ignored to improve readability of the

ASTRID tree topology. T3SS categories precede the name of the operon (e.g. “8 Pand7” is operon Pand7 belonging to category 8), downloaded from T3Enc database v1.0 (Hu et al. 2017). Tip labels for T3SS in the three *D. discoideum*-symbiont genomes and in *B. mallei* and *B. pseudomallei* are shown in bold font face with gene IDs for ease of reference. The clade containing the shared T3SS operon is shaded.

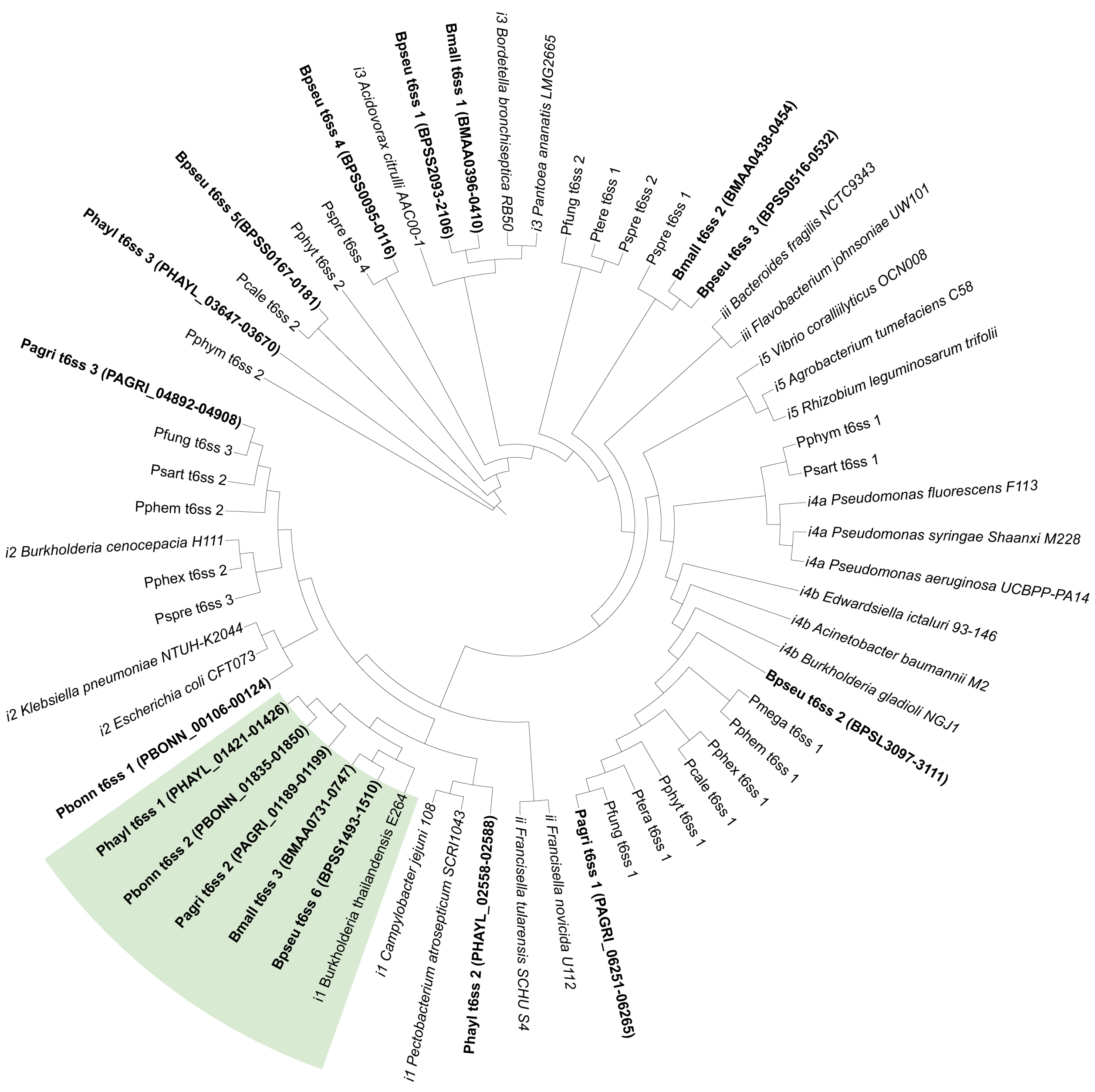

Figure S5. Type 6 secretion systems categorized using the conserved component genes tssB (sheath; COG3516), tssC (sheath; COG3517), and tssF (baseplate; COG3519). Branch lengths were ignored to improve readability of the ASTRID tree topology. T6SS categories precede the name of the strain to which the operon belongs (e.g. "ii Francisella novicida U112" is belongs to category ii), downloaded from SecReT6 databse v3.0 (Li et al. 2015). T6SS in the three *D. discoideum*-symbiont genomes and in *B. mallei* and *B. pseudomallei* are shown in bold font face with gene IDs for ease of reference. The clade containing the shared T6SS operon is shaded.
